# Evolutionary ecology of natural comammox *Nitrospira* populations

**DOI:** 10.1101/2020.09.24.311399

**Authors:** Alejandro Palomo, Arnaud Dechesne, Otto X. Cordero, Barth F. Smets

## Abstract

Microbes commonly exists in diverse and complex communities where species interact, and their genomic repertoires evolve over time. Our understanding of species interactions and evolution has increased in the last decades, but most studies of evolutionary dynamics are based on single species in isolation or in experimental systems composed of few interacting species. Here, we use the microbial ecosystem found in groundwater-fed sand filters as a model to avoid this limitation. In these open systems, diverse microbial communities experience relatively stable conditions, and the coupling between chemical and biological processes is generally well defined. Metagenomic analysis of 12 sand filters revealed systematic co-occurrence of at least five comammox *Nitrospira* species, likely promoted by low ammonium concentrations. These *Nitrospira* species showed intra-population sequence diversity, although possible clonal expansion was detected in few abundant local comammox populations. They showed low homologous recombination and strong purifying selection, the latter process being especially strong in genes essential in energy metabolism. Positive selection was detected on genes related to resistance to foreign DNA and phages. We found that, compared to other habitats, groundwater-fed sand filters impose strong purifying selection and low recombination on comammox *Nitrospira* populations. These results suggest that evolutionary processes are more affected by habitat type than by species identity. Together, this study improves our understanding of species interactions and evolution in complex microbial communities, and sheds light on the environmental dependency of evolutionary processes.

## Introduction

Microorganisms dominate the tree of life based on species diversity, and they play an essential role in all global biogeochemical cycles. Microbial species interact with each other and with the environment (ecological processes), and also undergo changes in their genomic repertoire over time (evolutionary processes). Yet, the interaction between ecological and evolutionary processes is largely unknown, especially for complex open communities. For many years, most studies of microbial communities in open, complex environments have focused on ecological aspects as it was believed that evolutionary changes happen at much larger timescale (1). However, in recent years, with the development of population-genomics analysis, researchers have started to investigate jointly the ecological and evolutionary processes. Yet, most studies of evolutionary dynamics remain based on single species in isolation (2) or on experimental systems composed of only a few interacting species (3). While these analyses have helped to better understand some aspects of evolutionary patterns, they have limitations because they lack many characteristics of true natural populations (e.g., spatial structure, existence of microdiversity, predation, immigration). On the other hand, observing populations in the wild also has limitations because the conditions vary with little control (hence, uncontrolled variation in population size, selection regime) and because the typically unknown ecophysiology of retrieved genomes makes it difficult to interpret the observed patterns. Therefore, studying well-defined model microbial ecosystems can help to understand ecological and evolutionary processes in microbial communities (4).

Rapid sand filters (RSF), widely used to produce drinking water from surface- or groundwater, are useful model systems as they are characterized by stable conditions; active growth, primarily driven by the oxidation of ammonia, methane, and other inorganic compounds present at low concentration in the influent water; large populations (10^9^ – 10^10^ cells/g); significant mixing; continuous but limited immigration from prokaryotes in the influent water; no dispersal between separate sand filters (resulting in allopatric populations); and relatively well defined coupling between chemical and biological processes (5–8). In addition, the microbial communities inhabiting these systems, which are usually stable across time (9), have been broadly described, showing a general dominance of complete ammonia oxidizers (comammox) (6, 10, 11). These recently discovered microorganisms are expected to have a relatively simple ecology (due to their chemolithoautotrophic metabolism) (12), yet are poorly studied in terms of what drives their diversity, distribution and evolution. Furthermore, as comammox bacteria occur in RSF as coexisting populations (10, 13), RSF offer an opportunity for resolving fine-scale genomic heterogeneity within closely related strains, and investigate if they show similar patterns in evolutionary processes (such as selection or recombination).

Of particular interest is to what extent the evolutionary processes that drive the diversification of comammox *Nitrospira* are dependent on their environment, as opposed to intrinsic properties of the species. Environmental dependency of microbial evolution has been investigated from different perspectives. Several studies have focused on genome signatures variations (GC, tetranucleotide signatures, codon usage, purine-pyrimidine ratio) associated with different environments (reviewed in Dutta and Paul (2012) (14)). Others have studied bacterial adaptation to shifting environments (15), or have targeted a specific evolutionary process across several lifestyles (e.g.: homologous recombination (16) or selection (17)). Most of these studies, however, considered different species living in different environments, or closely related species with a different lifestyle (i.e. free-living organisms vs pathogens). Yet, little is known about ongoing evolutionary processes of species belonging to the same lineage inhabiting different open environments. In this study, taking advantage of the multiple comammox species present in several groundwater-fed RSF, we thoroughly investigated evolutionary processes in this environment, and compared these observations with those in comammox species inhabiting other environments.

## Results and Discussion

We examined ecological and evolutionary patterns within comammox dominated-bacterial communities inhabiting groundwater-fed rapid sand filters. To that end, we retrieved metagenome-assembled genomes (MAGs) from 12 similarly operated waterworks in Denmark, using a combination of automatic and manual binning, followed by several refinement steps to improve the bin quality (Fig. S1). To remove redundancy, a single representative was selected for each set of genomes that shared an average nucleotide identity (ANI) greater than 99%. This resulted in a total of 189 MAGs (genome completeness = 83.9% ± 13.6, contamination = 1.9% ± 1.4; Table S1), 18 of them classified as *Nitrospira* spp. (completeness = 89.1% ± 9.2, contamination = 2.5% ± 1.1; Table S2). These *Nitrospira* MAGs spanned 16 putative species (further on simply referred to as ‘species’) using a threshold average nucleotide identity (ANI) of ≥ 95% (18–20). The phylogenomic analysis placed one *Nitrospira* species into lineage I, 14 into lineage II, and one into other lineages (Fig. 1). Of the 16 *Nitrospira* species, 12 were classified as comammox *Nitrospira* (5 clade A and 7 clade B) (Fig. 1). As expected, the genomes classified as comammox *Nitrospira* also contained genes essential for complete ammonia oxidation, such as those of the ammonia monooxygenase (AMO) and hydroxylamine dehydrogenase (HAO) operons. Further description of the *Nitrospira* MAGs can be found elsewhere (21).

**Fig. 1.**
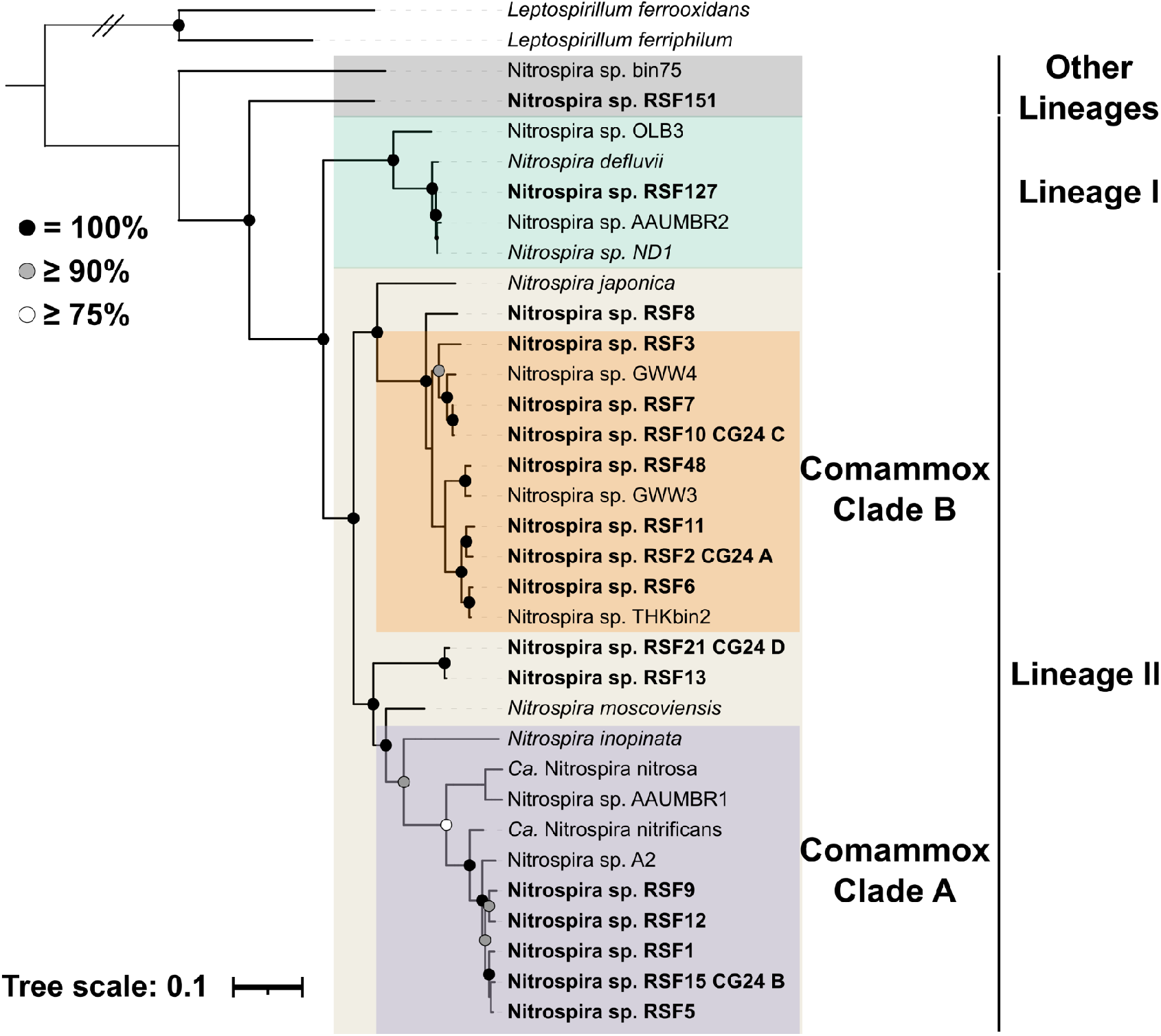
Phylogenomic affiliation of *Nitrospira* MAGs retrieved from 12 waterworks. A phylogenetic tree was built based on the concatenation of 120 proteins. *Nitrospira* MAGs retrieved in this study are highlighted in boldface. Lineages and sublineages are shown with colours (lineage I, green; lineage II, light brown; Other lineages, grey; comammox clade A purple; comammox clade B, orange). *Leptospirillum* was used to root the tree. The strength of support for internal nodes as assessed by bootstrap replicates is indicated as coloured circles (top left legend).

*Nitrospira* species comprised a large proportion of the microbial communities of the waterworks (27 - 70%), and comammox represented a large fraction of *Nitrospira* spp. (76 - 98%) (Fig. 2 and Table S3). For most of the RSFs, the variation among the abundance of the different *Nitrospira* MAGs between duplicate samples was low (Fig. S2), suggesting a high spatial homogeneity in the top layer of the filters, probably due to the frequent mixing caused by backwash, and hence supporting the representativity of our sampling. Multiple *Nitrospira* species (at least 5; 10 on average) co-occurred in all the waterworks (Fig. 2). However, there was no consistent dominance pattern. In some cases, a single species, but not always the same, dominated (WW4 and WW5), while, in other, two (WW9 and WW12) or more species dominated (Fig. 2). The chemical characteristics of the water explained 57% of the variance in *Nitrospira* composition (permutation test: p < 0.001; Table S4), suggesting that water chemistry is a strong filter for the assembly of these nitrifying communities. Among the measured water constituents, the influent ammonium concentration best explained *Nitrospira* distribution (explained 18%; permutation test: padjust = 0.02) (Table S4). Higher comammox species richness was detected in waterworks treating lower ammonium concentration (Fig. S3, R^2^ = 0.54, p < 0.01). Canonical *Nitrospira* and canonical ammonia oxidizers were more abundant in waterworks receiving influent with higher ammonium (Fig. S4). These observations are in line with the prediction that higher ammonium concentration favours the emergence of division of labour between canonical ammonia and nitrite oxidiser (22). Nevertheless, we observed that a comammox species (RSF3) seemed to cope with slightly higher ammonium concentrations as well (Fig. S4). Interesting, besides the mentioned RSF3, the other comammox *Nitrospira* spp. tended to cluster with those ones sharing clade affiliation (Fig. S4), which is based on the phylogeny of ammonia monooxygenase subunit. On another note, the distribution pattern of *Nitrospira* species across the waterworks was not related to their geographic distance (Mantel test: r statistics = 0.08 and significance > 0.05).

**Fig. 2.**
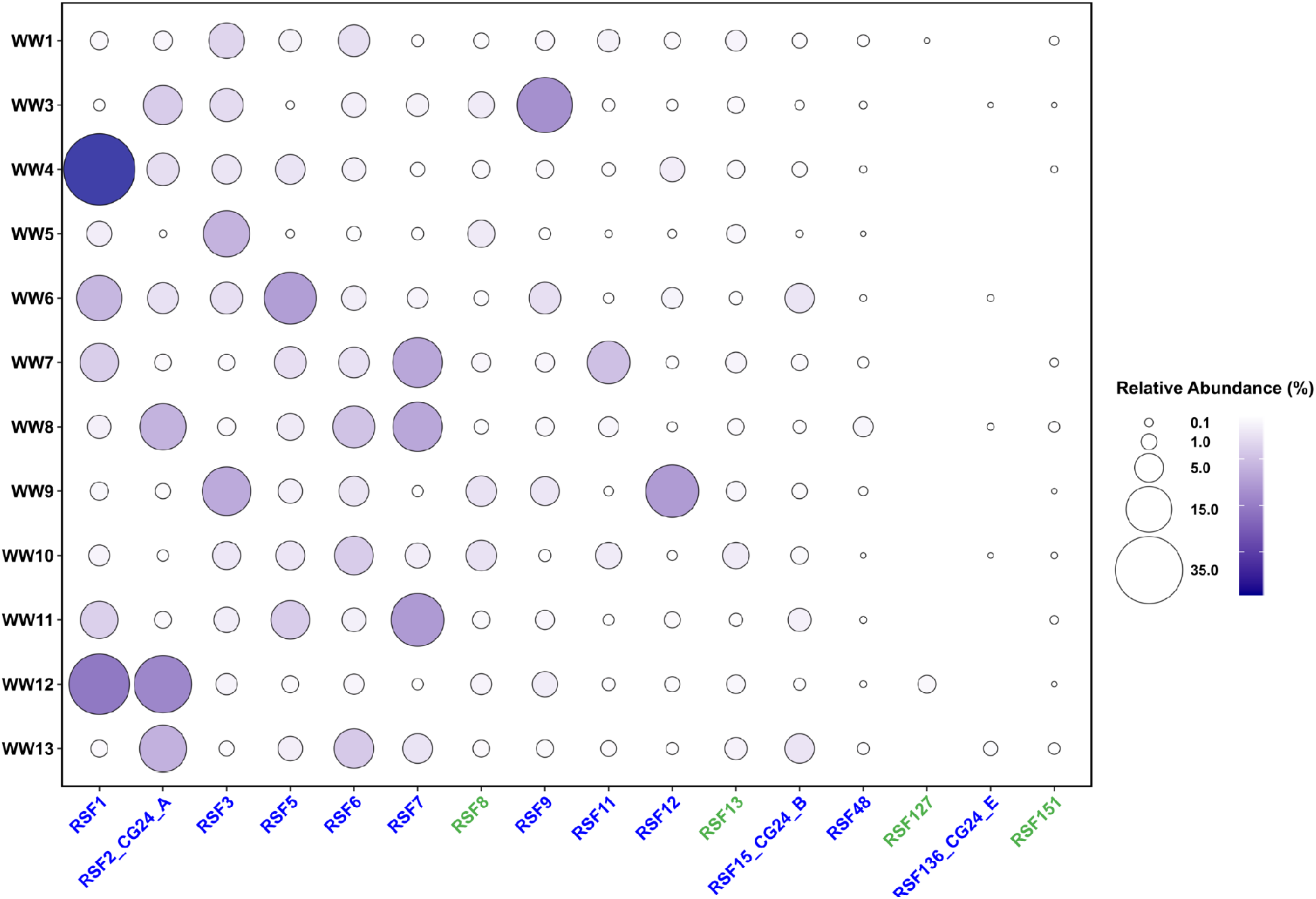
Abundance of *Nitrospira* species across 12 waterworks. Relative abundance of 16 *Nitrospira* species in 12 waterworks. Comammox and canonical *Nitrospira* species are denoted in blue and green, respectively.

Significant correlation among the abundances of the comammox species was rare; only few strong positive correlations were detected (RSF5 and RSF15_CG24_B (ρ=0.84), RSF6 and RSF11 (ρ=0.84)), while no significant negative correlations were observed (Fig. S5). This absence of obvious competitive exclusion pattern, together with the aforementioned description of co-existence of multiple comammox species, suggest that these microorganisms seem to exploit slightly different niches. A possible explanation might be that the co-occurrent comammox *Nitrospira* species had different ammonia affinities, and they would occupy different sites in the highly porous filter material (23). So far, little is known about the variability of ammonia affinity in comammox *Nitrospira* as this feature has only been reported for two comammox species (24, 25), however, a wide range of ammonia affinities have been observed in other ammonia oxidizers (26). Another reason could lie in the abundance of unique genes in each comammox *Nitrospira* species (> 250 in average (21)). Although the function of most of them is still unknown, these unique genes might promote ecological variation. Traits such as chemotactic strategies, attachment to particles strategies, secondary metabolism or defence against predation have been proposed to explain this phenomenon in other coexisting microorganisms (Reviewed in Louca *et al.*, (2018) (27)).

### Microdiversity within *Nitrospira* species

Strain-level analysis across the waterworks revealed that the *Nitrospira* populations contained intra-population sequence diversity. We exploited the shotgun metagenomic data to perform strain-level analyses on the 12 most abundant *Nitrospira* species (genome completeness = 92.4% ± 5.5, contamination = 2.7% ± 0.8) based on single nucleotide polymorphisms (SNPs). The number of SNPs/Mbp in the populations across the waterworks ranged from 14,437 to 45,664 (Table S5). Looking into the populations at local scale (species within waterworks), the number of SNPs/Mbp ranged from 249 to 37,663 (Table S6).

We observed a wide range of microdiversity (measured as nucleotide diversity (π)) among populations (Fig. 3A): canonical *Nitrospira* RSF8 was the most diverse species, with three times more nucleotide diversity than the less diverse *Nitrospira* species of our study (RSF1 and RSF12) (Fig. 3A). Depending on the *Nitrospira* population, we detected instances of both homogeneous microdiversity across the waterworks (e.g., RSF5 and RSF8), as well as a high microdiversity variation depending on the waterworks (e.g., RSF1, RSF9 and RSF11) (Fig. S6). Based on our observations of high species-level comammox *Nitrospira* diversity at low ammonium concentration, we hypothesised that such conditions also promote high microdiversity. However, this was not the case, as the correlations of microdiversity with ammonium concentration or with comammox species richness were not significant for any species (p > 0.05). This suggests different drivers for inter-species vs within-species diversities.

**Fig. 3.**
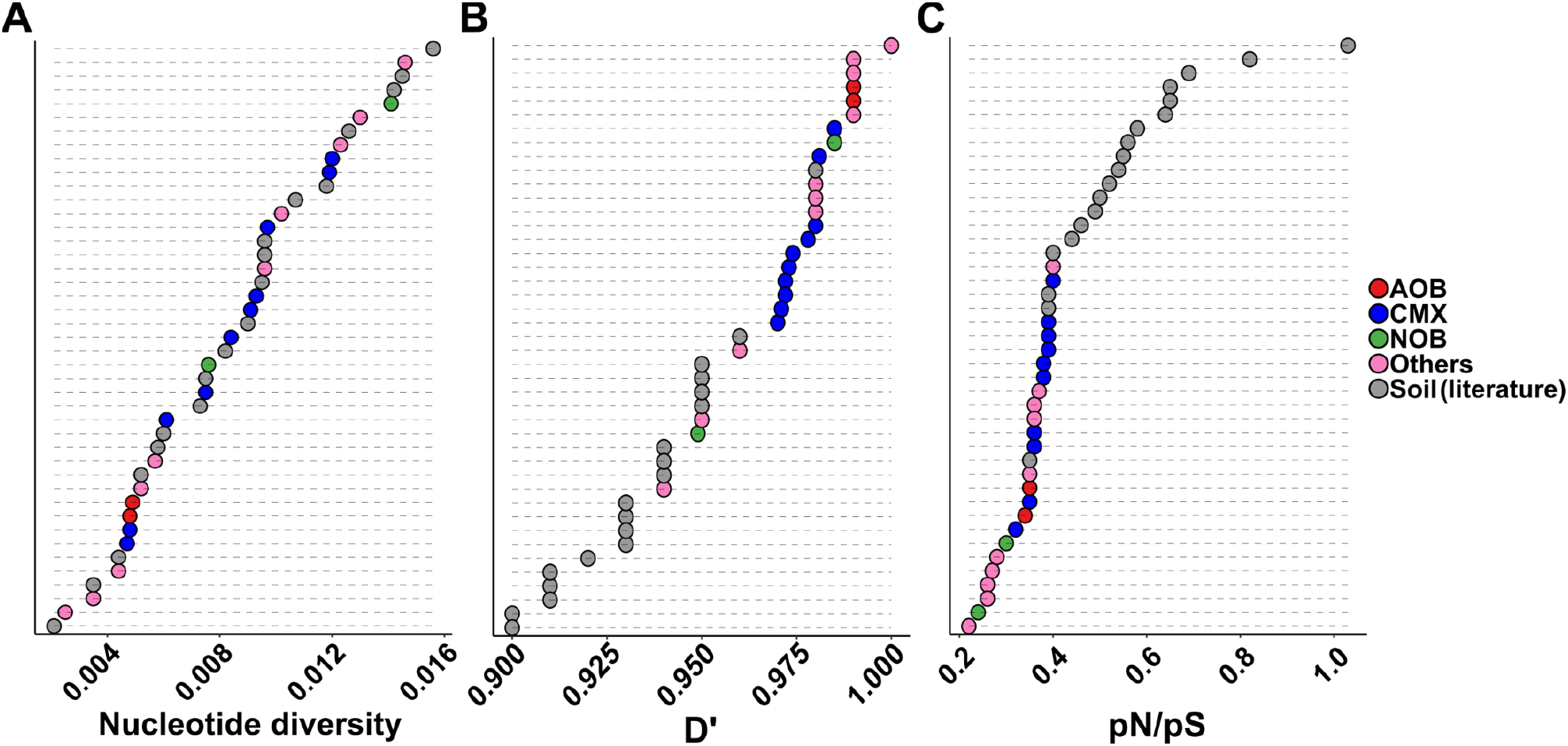
Evolutionary metrics of *Nitrospira* populations across 12 waterworks compared to other organisms and locations. A) Nucleotide diversity (π) of the most abundant bacterial populations across 12 waterworks, and the most abundant bacterial populations across a grassland meadow (denoted as ‘Soil’) [38]. B) Homologous recombination (D’) of most abundant bacterial populations across 12 waterworks, and most abundant bacterial populations across grassland meadow. C) Selection (pN/pS ratio) of most abundant bacterial populations across 12 waterworks, and most abundant bacterial populations across grassland meadow. AOB: ammonia oxidizing bacteria, CMX: comammox, NOB: nitrite oxidizing bacteria, Others: non-nitrifying abundant bacteria present in the waterworks. The data reported in this table (including the MAGs ID) can be found in Table S5.

While all species showed significant microdiversity across waterworks, we detected a few highly abundant comammox populations with almost no microdiversity at the local scale (e.g., comammox *Nitrospira* RSF1 and RSF12 in in WW4 and WW9, respectively; Fig. S7), which suggests local clonal expansions of these comammox populations. Along the same line, the analysis of major allele frequencies of common SNPs revealed that only a fraction of the within-species variants diversity is found locally. Thus, for specific species, we detected different within-species variants among the waterworks, in some cases, distinct single variants being dominants (e.g.: comammox RSF3 in WW10B vs WW5; Fig. S8). These results contrast with species level observations, where all the diversity was represented in each waterworks (as they all contain most of the *Nitrospira* spp.; Fig. 2).

Similar to what we observed at species level, there was no significant correlation between similarity in subspecies composition and the geographic distance of the waterworks, with the exceptions of *Nitrospira* sp. RSF2 (p < 0.01) and *Nitrospira* sp. RSF8 (p < 0.05) (Fig. S9). However, we observed an inter-waterworks organisation of the genetic structure at most loci across the studied genomes, indicating that the *Nitrospira* populations were more similar within than between waterworks. For each gene, we calculated pairwise fixation indexes (F_*ST*_) to measure differences in allele frequencies between populations of the same species found in two distinct waterworks. The mean gene F_*ST*_ values were ≥ 15% for all *Nitrospira* species (Fig. S10), the most extreme case being RSF5 (F_ST_ > 40%) (Fig. S10). In few species (RSF2, RSF7 and RSF8), a higher dispersal (F_ST_ < 20%) of most alleles between waterworks was observed (Fig. S10). These observations differ from those on soil bacterial populations across a meadow obtained using the same approach, where most of the populations had mean gene F*_ST_* values < 5% (28). These contrasting results support the notion that populations in the waterworks are much more allopatric than the ones in other environments such as the mentioned meadow.

We also investigated local regions of the *Nitrospira* genomes with significantly higher F_*ST*_ values, as this is characteristic of local population-specific (here, in each waterworks) selective pressures (28). A few loci with unusually high F_*ST*_ were found in some of the *Nitrospira* populations (Fig. S11 and Table S7), one of them containing genes involved in nitrogen assimilation (*Nitrospira* sp. RSF2) (Table S7). However, only two of these loci with unusually high site-specific differentiation of alleles (high F_*ST*_) also had few recombinant events, and low nucleotide diversity (Table S7), which can be considered as a signal of recent selective sweep (28). These results suggest that, contrary to what has been observed in several natural populations (28–31), gene-specific sweeps seem to play a minor role in the evolution of *Nitrospira* spp. inhabiting the waterworks. A possible explanation could be the low recombination rate that characterised the waterworks *Nitrospira* populations (Fig. 3B, discussed below), as opposed to genome-wide sweeps that are associated to low recombination rates, gene-specific sweeps are expected to occur with high recombination rates (32).

Overall, across the 12 waterworks, all species presented significant genomic microdiversity but this diversity was not always represented locally, with a few occurrences of patterns consistent with clonal expansion. The reason for the difference of within-species diversity across waterworks is unknown but the allopatric nature of the communities likely contributes to their persistence.

### Evolutionary processes at whole-genome level

The *Nitrospira* populations were characterised by a low degree of homologous recombination. We investigated this evolutionary process based on linkage disequilibrium (D’ is only < 1 if all possible combinations of a pair of biallelic sites are observed (33); lower D’ values indicate higher levels of homologous recombination; Fig. 3B). A similar low degree of homologous recombination was observed for other non-*Nitrospira* abundant populations inhabiting the waterworks (n=12; genome completeness = 92.6% ± 5.5, contamination = 2.1% ± 2.0) (Fig. 3B and Table S5). In general, this evolutionary process was lower in the waterworks populations than in populations inhabiting a grassland meadow (Fig. 3B), where a similar strain-level analysis was conducted (28). To further examine the relative effect of homologous recombination on the genetic diversification of the populations, we measured the rates at which nucleotides become substituted as a result of recombination versus mutation using the *r/m* ratio. Most of the *Nitrospira* populations had a relatively low *r/m* (*r/m* < 2) compared to recombinogenic species reported in literature (*r/m* > 4) (34) (Fig. S12), although in one case (RSF15_CG24_B) the rate was similar to the value reported for a *S. flavogriseus* population (*r/m* = 28) considered to be approaching panmixia (35). Overall, these results suggest a low effect of recombination in the populations inhabiting the waterworks. Increasing recombination rate has been associated with fluctuating environments as a source of variation which can accelerate adaptation favouring survival in this type of environments (36, 37). On the other hand, constant environments - as the waterworks studied here - tend to reduce the recombination rate of their residents (36).

The *Nitrospira* populations were characterised by strong purifying selection. We used the relation between non-synonymous and synonymous polymorphisms (pN/pS) to investigate this evolutionary process. We detected pN/pS < 1, indicating purifying selection, for all *Nitrospira* species (Fig. 3C). Similar results were observed for other abundant populations of the waterworks (Fig. 3C). Purifying selection has frequently been observed in wild populations, and it was the case for populations inhabiting a grassland meadow (28) (pN/pS = 0.56 ± 0.17, n = 19), but this process seems to be especially strong in the waterworks populations (pN/pS = 0.34 ± 0.05, n = 24) (Two-Sample t-test, p < 0.0001) (Fig. 3C). This suggests that their genomes could have reached an adaptive optimum for this stable environment, which is maintained by purging non-synonymous mutations.

Interestingly, the degree of recombination and diversity across different *Nitrospira* populations varied substantially with habitat (Fig. 4). High variability of recombination in closely related bacterial species has occasionally been reported (38), and lifestyle appears as one of the most relevant factors to explain this variability (16, 38). Our analysis in *Nitrospira* populations from different habitats (drinking water treatment plants (DWTP), freshwaters and soils) suggests that the environment also influences ongoing evolutionary processes: different bacterial types in the same environment tended to share similar features (Fig. 4), while the evolutionary characteristics of comammox *Nitrospira* populations differed depending on the environment where they were retrieved (Fig. 4). Comammox species in the studied waterworks and in other DWTP were characterised by low recombination, strong purifying selection and moderate microdiversity (Fig. 4). On the other hand, comammox present in freshwater and in soils had higher microdiversity and, especially, recombination rate (Fig. 4 and Table S5). Intriguingly, we consistently observed that, in drinking water treatment systems, canonical *Nitrospira* species showed features similar to those of comammox *Nitrospira* but with even stronger purifying selection (Fig. 4). This feature, together with the much lower richness observed in canonical *Nitrospira* compared to comammox bacteria (Fig. 2), suggests that competition can play a more intense role in canonical *Nitrospira*, which might select for few species optimally adapted to this type of stable environment. However, a broader analysis is required to confirm this hypothesis.

**Fig. 4.**
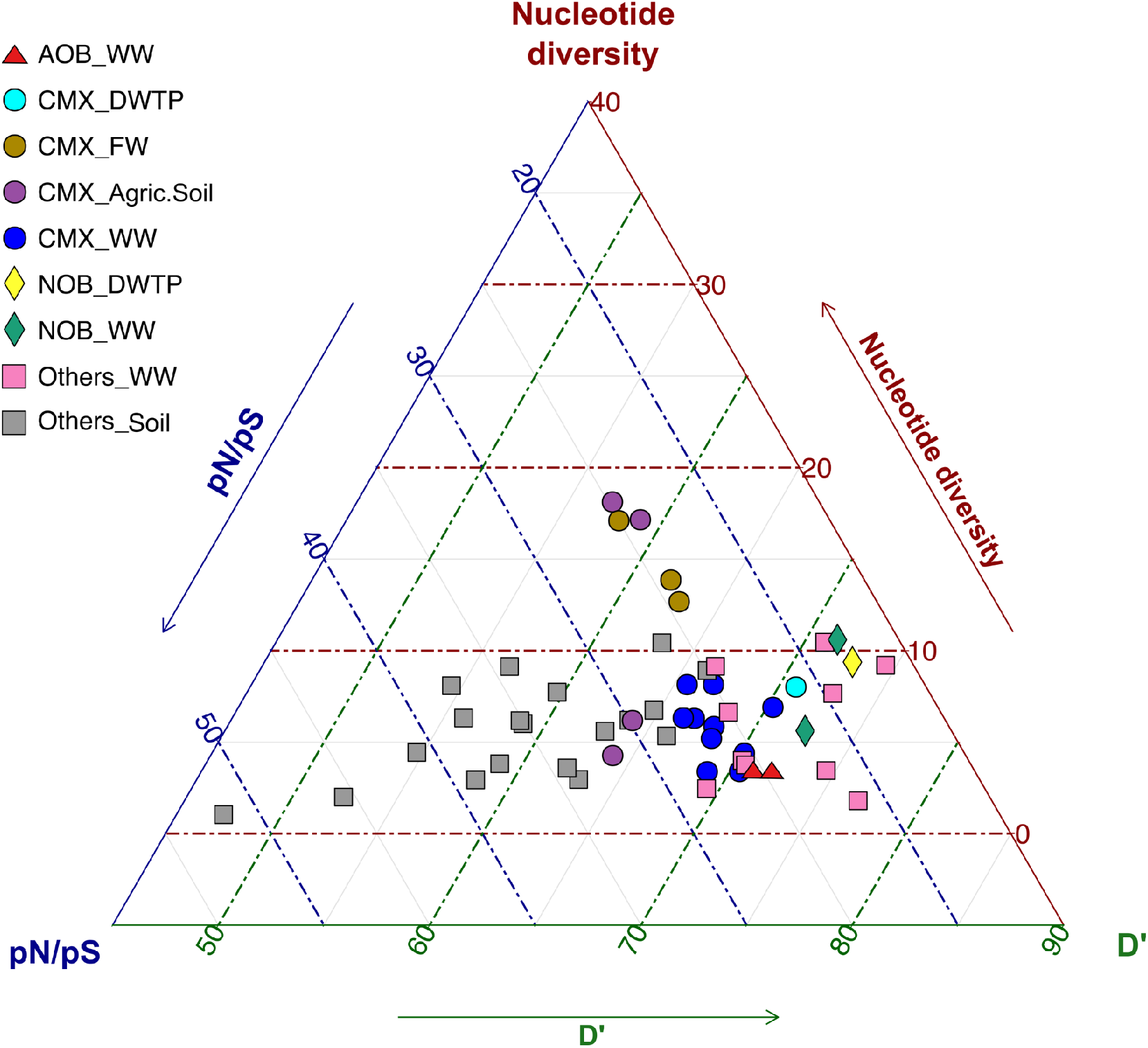
Impact of environment and microbial type in evolutionary metrics. Triplot composed of the nucleotide diversity, pN/pS ratio and D’ values for the bacterial populations of this study (WW), the most abundant bacterial populations across a grassland meadow [38] (Soil), and other abundant *Nitrospira* populations recovered from other systems (Table S5; CMX_FW, CMX_DWTP, NOB_DWTP, and CMX_Agric.Soil). The data reported in this table (including the MAGs ID) can be found in Table S5.

### Evolutionary processes at the gene level

In addition to a genome-wide analysis, we investigated the evolutionary processes at the gene level. In the studied *Nitrospira* populations, genes involved in nitrification (ammonia monooxygenase: *amoA* and *amoB*; hydroxylamine dehydrogenase: *haoA* and *haoB*; nitrite oxidoreductase: *nxrA* and *nxrB*) generally had a similar nucleotide diversity (π) (Fig. 5A) and homologous recombination rate (D’) (Fig. 5B) compared to the rest of the genome, but with higher levels of purifying selection (pN/pS) (Fig. 5C). The nucleotide diversities of genes related to nitrification were very similar with the exception of *amoB*, which had a significantly lower nucleotide diversity than *nxrB* (p < 0.05) (Fig. 5A). A similar pattern was detected for the recombination, but in this case *amoA*, as well as *amoB*, had significantly lower recombination than *nxrB* (p < 0.05) (Fig. 5B). We observed a very strong purifying selection for most of the nitrifying genes, especially for *amoA*, *nxrA*, and *nxrB* (p < 0.01) (Fig. 5C). In the case of *nxrB*, not a single non-synonymous mutation was found in most of the *Nitrospira* species (0-1 non-synonymous sites vs 17-66 synonymous sites), even though this gene had a higher nucleotide diversity and homologous recombination (Fig. 5A and Fig. 5B). Our observations on selection are in line with previous studies, as generally, essential genes and enzymes catalysing reactions that are difficult to by-pass through alternative pathways are subject to higher purifying selection compared to nonessential ones (39–42).

**Fig. 5.**
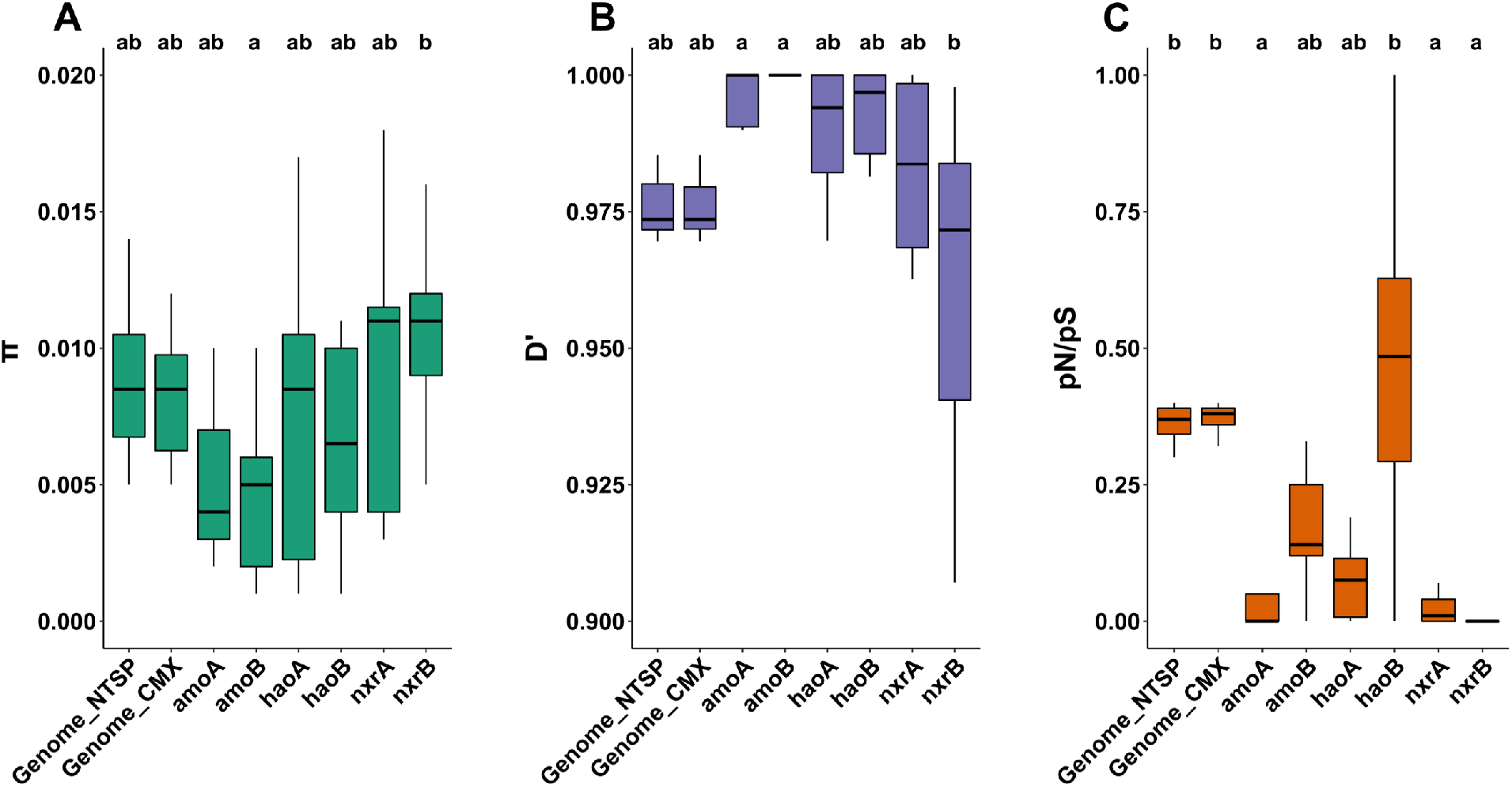
Evolutionary metrics of nitrification genes in *Nitrospira* populations across 12 waterworks. Left) Boxplot of nucleotide diversity of *Nitrospira* bacterial populations for whole genome (all *Nitrospira* and comammox *Nitrospira*) and nitrification genes. Differences between the mean nucleotide diversities were assessed by a Dunn’s test; same letter means no significant difference (p < 0.05). Middle) Boxplot of linkage disequilibrium of *Nitrospira* bacterial populations for whole genome (all *Nitrospira* and comammox *Nitrospira*) and nitrification genes. Differences between the mean linkage disequilibriums were assessed by a Dunn’s test; same letter means no significant difference (p < 0.05). Right) Boxplot of pN/pS ratios of *Nitrospira* bacterial populations for whole genome (all *Nitrospira* and comammox *Nitrospira*) and nitrification genes. Differences between the mean pN/pS ratios were assessed by a Dunn’s test; same letter means no significant difference (p < 0.01).

Even though the average pN/pS values were below 1 in all *Nitrospira* species (Fig. 3), indicating purifying selection, genes with pN/pS values above 1 and significantly higher than the genomic average were detected in each species (Fig. S13). Many of those genes under positive selection were related to defence mechanism against phages (e.g. genes putatively involved in phage entry into cells, ribonucleases, genes coding proteins associated to restriction-modification systems, and genes related to toxin-antitoxin systems (Table S8)). Comparable findings were made in other abundant species from the waterworks (Table S9), in the additional *Nitrospira* populations retrieved from other environments (Table S9), as well as in *E. coli* and *Vibrio* sp. strains, respectively (43, 44). These observations suggest that positive selection in phage-related genes is widespread across bacteria, and highlights the evolutionary arms race occurring between phages and bacteria as an important driver in bacterial ecology and evolution (32, 45, 46). Additionally, we found nondefense, mobile genetic elements, such as transposons and integrases, with significantly higher pN/pS values than the genome average in the *Nitrospira* spp. (Table S10).

## Conclusions

A major unresolved question is how the relationship between ecology and evolution shapes complex communities in the environments. Here we use a model microbial system with relatively stable conditions to examine this question. By analysing the microbial communities in 12 groundwater-fed rapid sand filters, we observed that co-occurring comammox *Nitrospira* spp. dominate all sites, but with local differences in species composition. This suggest that the abundant comammox *Nitrospira* spp. exploit slightly different niches, which could partially be explained by the inlet water chemistry composition and the different genetic repertory. At subspecies level, comammox *Nitrospira* spp. are characterised by strong purifying selection and low recombination. These features, together with the occasional genome sweeps we detected, suggest that the subspecies possibly occupy a narrow niche to which they are optimally adapted. In contrast, we also showed that the relative magnitude of these evolutionary processes was different in comammox *Nitrospira* in habitats where environmental conditions are less stable and immigration more intense. Thus, we conclude that the evolutionary processes that drive the diversification of *Nitrospira* are dependent on the environment, as opposed to intrinsic properties of the species.

## Materials and methods

### Sampling, sequencing, and metagenomic assembled genomes recovery

The sampling description, DNA extraction and sequencing have been previously described (21). Briefly, filter material was collected from two locations at the top of the filters of 12 Danish waterworks. DNA was extracted from 0.5 g of sand material using the MP FastDNA Spin Kit (MP Biomedicals LLC, Solon, USA). DNA libraries were generated using the 24 extracted DNA with the Nextera XT DNA library preparation kit (Illumina Inc.) according to the manufacturer’s instructions. Samples were sequenced in one lane, with 2×150 paired read sequencing on the Illumina HiSeq4000 at BGI’s facility in Copenhagen. Generated reads were trimmed and filtered, including adapters removal, using Trimmomatic v0.22 (threshold quality=15; minimum length=40) (47). FastQC (Babraham Bioinformatics (http://www.bioinformatics.babraham.ac.uk/projects/fastqc/)) was used to evaluate the quality of the obtained reads. Co-assembly of high-quality reads was conducted using IDBA-UD (48) with the options --pre_correction --min_contig 1000. Additionally, 24 single-sample assemblies were performed following the same procedure. As it has been shown that the combination of multiple binning algorithms outperforms the usage of one single algorithm (49), here we performed metagenomic binning using MetaBAT (50), MaxBin2 (51), CONCOCT (52), MyCC (53), Binsanity (54) and COCACOLA (55) (Fig. S1). The quality of the resulted genomes was improved using Binning refiner (56). Simultaneously, we ran the software metaWRAP (57), which also take advantage of multiple binning tools to recover genomes from the co-assembly. The generated bins were improved using the refinement module of metaWRAP. Selection of the best genomes recovered with the different binning algorithms was done with DAS Tool (58). dRep (59) was applied to dereplicate all the selected best bins with the secondary clustering threshold set at 99% genome-wide Average Nucleotide Identity (gANI). On the other hand, mmgenome (60) was applied to the single-sample assemblies to recover *Nitrospira* genomes following the strategy described elsewhere (13). The generated *Nitrospira* genomes were combined with the dereplicated bins, and subjected to the reassembly module of metaWRAP with the aim of improving the quality of the bins. dRep at 99% gANI was used again for dereplication. RefineM (--cov_corr = 0.8) (61) was used to refine the resulting dereplicated bins by removing scaffolds with divergent GC-content or tetranucleotide frequencies. Furthermore, the binning and refinement modules from metaWRAP were applied to the co-assembly of the six RSF samples used in Palomo et al. (2018) (5). Obtained bins together with the reported ones in Palomo et al. (2016 and 2018) (5, 13) were dereplicated using dRep at 99% gANI. A refinement step with RefineM was applied on the resulting dereplicated bins. These refined bins, together with the bins recovered from the 12 waterworks of this study were dereplicated as described above. In addition, the assembly quality of the *Nitrospira* bins were improved by alignment against related complete or draft genomes using the Multi-Draft based Scaffolder (MeDuSa) (62). The overall procedure here described can be visualised in the Fig. S1. The completeness and contamination of the bins were evaluated using CheckM (63).

### Species abundance estimation

A 95% average nucleotide identity (ANI) cut-off was used to define species as proposed by Klappenbach *et al*. (2007) (18). The retrieved MAGs were dereplicated using dRep with the secondary clustering threshold set at 95% gANI. Among the genomes classified as belonging to the same species, the one with higher quality was chosen as representative genome. The species abundance and coverage of each representative genome across the metagenomes were assessed using MIDAS (64). Briefly, MIDAS uses reads mapped to 15 universal single-copy gene families (with ability to accurately recruit metagenomic reads to the correct species (64)) to estimate the abundance and coverage of bacterial species from a shotgun metagenome. We used the species retrieved in this study to build the database of universal-single-copy genes.

### Genome classification and annotation

MAGs were classified using the classify workflow of the GTDB-Tk v.0.1.3 tool (65). Open reading frames were predicted using Prodigal v. 2.63 (66), and annotated using blastp (67) against NCBI nr (68), UniProt (69), KEGG (70), PFAM (71) and eggNOG (72). Genes were assigned to antiphage defense systems using the strategy described in Doron *et al.* (2018) (73).

### Phylogenetic analysis

Phylogenetic analyses of *Nitrospira* genomes were conducted with the GTDB-Tk v.0.1.3 tool (65) [21] using the *de novo* workflow with a set of 120 single copy marker proteins and the genome taxonomy database (GTDB) (74). Concatenated alignments were used to construct a maximum likelihood tree using RAxML v. 8.2.11 (75) with 200 rapid bootstraps (determined using the autoMRE option) and the LG likelihood model of amino acid substitution with gamma distributed rates and fraction of invariant sites (-m PROTGAMMAILGF; best model determined using ProtTest v. 3.4.2 (76)). The tree was rooted using two *Leptospirillum* species as outgroup. The rooted tree was visualized using the online web tool from the Interactive Tree of Life (iTol) (77).

### Read mapping, SNP calling, and population genomic analysis

The population genomic analysis was done following the approach described in Crits-Christoph et al (2020) (28). High-quality reads were mapped to an indexed database of the 176 species MAGs recovered from the waterworks using BWA-MEM (78). The resulting alignments were filtered using samtools (79) view -q30 to remove reads with mapping quality less than 30, and also with the script filter_reads.py (28) (with the options: -m 96 to retain reads with a percent identity of at least 96% to the reference; and -q 2 to assure uniquely best mapping read pairs in the index). Downstream population genomic analysis was performed on the 24 most abundant species genomes (12 *Nitrospira* ones, and 12 other abundant species genomes). For each of these species genomes, we analysed data in samples that passed a cutoff of at least 50% of the genome being covered with at least 5× coverage. 149 out of 576 sample genome comparisons (24 genomes × 24 samples) passed this minimum requirement. Sample read mappings were pooled by each waterworks and by all samples across the waterworks. Nucleotide diversity (π), linkage disequilibrium (D’) and pN/pS ratio were calculated for each sample, each waterworks and across all the waterworks as described elsewhere (28) using the scripts provided by the authors. F_*ST*_ was calculated following the same procedure but on sites segregating across two waterworks being compared (for all the possible waterworks comparisons). As Crits-Christoph *et al*. (2020) (28) recommended, only sites with a coverage of at least 20× in each waterworks was used to calculate F_*ST*_. In addition, genes with coverage in a waterworks outside of the range of two standard deviations were excluded from the analysis. As previously suggested (28), a two-sample Wilcoxon test was conducted to find out if average linkage of highly differentiated loci differed from the genomic average for each species. Similarly, a two-sample *t*-tests was used to conclude if average nucleotide diversity of highly differentiated loci differed from the genomic average. Both sets of tests were corrected for multiple hypotheses using the Benjamini–Hochberg method. Recombination to mutation ratios were inferred using mcorr (34).

Same strain-level analysis as the one described above was conducted in other *Nitrospira* MAGs previously recovered (21) that passed a cutoff of at least 50% of the genome being covered with at least 5× coverage in any of the metagenomes were *Nitrospira* MAGs were found to be present (21).

### Statistical Analyses

All statistical tests were performed using R v3.5.2 (80). Due to the compositional nature of sequencing data (81), for all statistical analyses, species abundances were analysed as followed: zeros were replaced with an estimate value using the Count Zero Multiplicative approach with the zCompositions R package (82), and data were further centred log-ratio transformed. *Nitrospira* community dissimilarities were calculated using the Jaccard index. The correlation between the *Nitrospira* community dissimilarities and geographic distances was calculated using the Mantel test (significance obtained after 100,000 permutations). Same analysis was used to assess the correlation between the *Nitrospira* community dissimilarities and the water composition dissimilarity, as well as the correlation between major allele dissimilarities and geographic distances.

Proportionality between abundances of the species across the 24 metagenomes were calculated using the propr R package (83) (with the options metric = “rho”, ivar = “clr”) and visualised using the corrplot R package (84). For the network analysis, the function getNetwork from propr R package was used to retain proportionalities > 0.56 (FDR < 5%). The network was visualised using the igraph R package (85).

Redundancy analysis (RDA) was performed in a stepwise way using the *ordiR2step* function of the *vegan* package (86). The analysis was conducted using centred log-ratio transformed *Nitrospira* species abundances and chemical data of influent water. The constrained ordination model and the variable significance were determined by permutation tests (1000 permutations) with anova.cca in vegan. Before this analysis, collinearity among explanatory variables was evaluated using variance inflation factor (VIF) in the *fmsb* package (87). The function *vif_func* was used to perform a stepwise approach until all variables above a VIF of 5 were excluded (88). Ternary plot was performed using the R package ggtern (89) using the nucleotide diversity, pN/pS ratio and D’ values for the bacterial populations retrieved from the waterworks, as well as most abundant bacterial populations across grassland meadow (28), and other *Nitrospira* populations abundant in other systems (Table S5).

Differences between the mean nucleotide diversities of the nitrifying genes, whole *Nitrospira* genomes, and whole comammox *Nitrospira* genomes were assessed using Kruskal–Wallis ANOVA followed by Dunn’s test with the Holm-Bonferroni correction. Same analysis was performed for linkage disequilibrium and pN/pS ratios.

### Chemical analysis of influent water

Ammonium was measured using a standard colorimetric salicylate and hypochlorite method (90), while nitrite was analysed using a standard method adapted from Grasshoff *et al*. (1983) (91). Nitrate and sulfate were measured by ion chromatography according to AWWA-WEF method 4110 (92). Iron, manganese and copper were determined by ICP-MS (7700x, Agilent Technologies), while calcium by ICP-OES (Varian, Vista-MPX CCD Simultaneous ICP-OES). Dissolved oxygen and pH were measured with a handheld meter (WTW, Multi 3430, with FDO^®^ 925 and SenTix^®^ 940 probes).

## Data availability

All raw sequence data and *Nitrospira* genomes retrieved from the Danish rapid sand filters have been deposited at the NCBI BioProject database under accession number PRJNA384587. The rest of the retrieved draft genomes from the Danish rapid sand filters are available on figshare (https://doi.org/10.6084/m9.figshare.12962075). Data produced from the strain-level analysis (nucleotide diversity, SPNs, linkage statistics and F_*ST*_ metrics) are available on figshare (https://figshare.com/projects/Evolutionary_ecology_of_natural_comammox_Nitrospira_populations/91217).

## Acknowledgements

We thank Gabriel E. Leventhal for helpful discussions on data analysis and theory. This research was supported by a research grant (13391, Expa-N) from VILLUM FONDEN.

## Authors contributions

A.P conceived the study and performed the bioinformatic analyses. A.P and A.D led interpretation of the results supported by O.X.C and B.F.S. A.P drafted the manuscript with input from A.D, O.X.C and B.F.S. All authors contributed to manuscript revision, and approved the final version of the manuscript.

## References

1. Hairston NG, Ellner SP, Geber MA, Yoshida T, Fox JA. 2005. Rapid evolution and the convergence of ecological and evolutionary time. Ecol Lett 8:1114–1127.

2. Barroso-Batista J, Sousa A, Lourenço M, Bergman M-L, Sobral D, Demengeot J, Xavier KB, Gordo I. 2014. The First Steps of Adaptation of Escherichia coli to the Gut Are Dominated by Soft Sweeps. PLoS Genet 10:e1004182.

3. Lawrence D, Fiegna F, Behrends V, Bundy JG, Phillimore AB, Bell T, Barraclough TG. 2012. Species Interactions Alter Evolutionary Responses to a Novel Environment. PLoS Biol 10:e1001330.

4. Denef VJ, Mueller RS, Banfield JF. 2010. AMD biofilms: using model communities to study microbial evolution and ecological complexity in nature. ISME J 4:599–610.

5. Palomo A, Jane Fowler S, Gülay A, Rasmussen S, Sicheritz-Ponten T, Smets BF. 2016. Metagenomic analysis of rapid gravity sand filter microbial communities suggests novel physiology of Nitrospira spp. ISME J 10:2569–2581.

6. Tatari K, Musovic S, Gülay A, Dechesne A, Albrechtsen H-J, Smets BF. 2017. Density and distribution of nitrifying guilds in rapid sand filters for drinking water production: Dominance of Nitrospira spp. Water Res 127:239–248.

7. Gülay A, Fowler SJ, Tatari K, Thamdrup B, Albrechtsen H-J, Al-Soud WA, Sørensen SJ, Smets BF. 2019. DNA- and RNA-SIP Reveal Nitrospira spp. as Key Drivers of Nitrification in Groundwater-Fed Biofilters. MBio 10.

8. Hu W, Liang J, Ju F, Wang Q, Liu R, Bai Y, Liu H, Qu J. 2020. Metagenomics Unravels Differential Microbiome Composition and Metabolic Potential in Rapid Sand Filters Purifying Surface Water Versus Groundwater. Environ Sci Technol 54:5197–5206.

9. Wagner FB, Diwan V, Dechesne A, Fowler SJ, Smets BF, Albrechtsen H-J. 2019. Copper-Induced Stimulation of Nitrification in Biological Rapid Sand Filters for Drinking Water Production by Proliferation of Nitrosomonas spp. Environ Sci Technol 53:12433–12441.

10. Fowler SJ, Palomo A, Dechesne A, Mines PD, Smets BF. 2018. Comammox Nitrospira are abundant ammonia oxidizers in diverse groundwater-fed rapid sand filter communities. Environ Microbiol 20:1002–1015.

11. Poghosyan L, Koch H, Frank J, van Kessel MAHJ, Cremers G, van Alen T, Jetten MSM, Op den Camp HJM, Lücker S. 2020. Metagenomic profiling of ammonia- and methane-oxidizing microorganisms in two sequential rapid sand filters. Water Res 185:116288.

12. Koch H, Kessel MAHJ van, Lücker S. 2018. Complete nitrification: insights into the ecophysiology of comammox Nitrospira. Appl Microbiol Biotechnol 1–13.

13. Palomo A, Pedersen AG, Fowler SJ, Dechesne A, Sicheritz-Pontén T, Smets BF. 2018. Comparative genomics sheds light on niche differentiation and the evolutionary history of comammox Nitrospira. ISME J 12:1779–1793.

14. Dutta C, Paul S. 2012. Microbial Lifestyle and Genome Signatures. Curr Genomics 13:153–162.

15. Scheuerl T, Hopkins M, Nowell RW, Rivett DW, Barraclough TG, Bell T. 2020. Bacterial adaptation is constrained in complex communities. Nat Commun 11:754.

16. González-Torres P, Rodríguez-Mateos F, Antón J, Gabaldón T. 2019. Impact of Homologous Recombination on the Evolution of Prokaryotic Core Genomes. MBio 10.

17. Martinez JL. 2009. The role of natural environments in the evolution of resistance traits in pathogenic bacteria. Proc R Soc B Biol Sci 276:2521–2530.

18. Klappenbach JA, Goris J, Vandamme P, Coenye T, Konstantinidis KT, Tiedje JM. 2007. DNA–DNA hybridization values and their relationship to whole-genome sequence similarities. Int J Syst Evol Microbiol 57:81–91.

19. Jain C, Rodriguez-R LM, Phillippy AM, Konstantinidis KT, Aluru S. 2018. High throughput ANI analysis of 90K prokaryotic genomes reveals clear species boundaries. Nat Commun 9:5114.

20. Olm MR, Crits-Christoph A, Diamond S, Lavy A, Matheus Carnevali PB, Banfield JF. 2020. Consistent Metagenome-Derived Metrics Verify and Delineate Bacterial Species Boundaries. mSystems 5.

21. Palomo A, Dechesne A, Smets BF. 2019. Genomic profiling of Nitrospira species reveals ecological success of comammox Nitrospira. bioRxiv https://doi.org/10.1101/612226.

22. Costa E, Pérez J, Kreft J-U. 2006. Why is metabolic labour divided in nitrification? Trends Microbiol 14:213–219.

23. Gülay A, Tatari K, Musovic S, Mateiu R V., Albrechtsen HJ, Smets BF. 2014. Internal porosity of mineral coating supports microbial activity in rapid sand filters for groundwater treatment. Appl Environ Microbiol 80:7010–7020.

24. Daims H, Lebedeva EV, Pjevac P, Han P, Herbold C, Albertsen M, Jehmlich N, Palatinszky M, Vierheilig J, Bulaev A, Kirkegaard RH, von Bergen M, Rattei T, Bendinger B, Nielsen PH, Wagner M. 2015. Complete nitrification by Nitrospira bacteria. Nature 528:504–509.

25. Sakoula D, Koch H, Frank J, Jetten MSM, van Kessel MAHJ, Lücker S. 2021. Enrichment and physiological characterization of a novel comammox Nitrospira indicates ammonium inhibition of complete nitrification. ISME J 15:1010–1024.

26. Jung M-Y, Sedlacek CJ, Kits KD, Mueller AJ, Rhee S-K, Hink L, Nicol GW, Bayer B, Lehtovirta-Morley L, Wright C, de la Torre JR, Herbold CW, Pjevac P, Daims H, Wagner M. 2021. Ammonia-oxidizing archaea possess a wide range of cellular ammonia affinities. ISME J https://doi.org/10.1038/s41396-021-01064-z.

27. Louca S, Polz MF, Mazel F, Albright MBN, Huber JA, O’Connor MI, Ackermann M, Hahn AS, Srivastava DS, Crowe SA, Doebeli M, Parfrey LW. 2018. Function and functional redundancy in microbial systems. Nat Ecol Evol https://doi.org/10.1038/s41559-018-0519-1.

28. Crits-Christoph A, Olm MR, Diamond S, Bouma-Gregson K, Banfield JF. 2020. Soil bacterial populations are shaped by recombination and gene-specific selection across a grassland meadow. ISME J 1–25.

29. Shapiro BJ, Friedman J, Cordero OX, Preheim SP, Timberlake SC, Szabó G, Polz MF, Alm EJ. 2012. Population Genomics of Early Events in the Ecological Differentiation of Bacteria. Science (80-) 336:48–51.

30. Rosen MJ, Davison M, Bhaya D, Fisher DS. 2015. Fine-scale diversity and extensive recombination in a quasisexual bacterial population occupying a broad niche. Science (80-) 348:1019–1023.

31. Bendall ML, Stevens SLR, Chan L, Malfatti S, Schwientek P, Tremblay J, Schackwitz W, Martin J, Pati A, Bushnell B, Froula J, Kang D, Tringe SG, Bertilsson S, Moran MA, Shade A, Newton RJ, Mcmahon KD, Malmstrom RR. 2016. Genome-wide selective sweeps and gene-specific sweeps in natural bacterial populations. Isme J 10:1589–1601.

32. Shapiro BJ, Leducq J-B, Mallet J. 2016. What Is Speciation? PLOS Genet 12:e1005860.

33. VanLiere JM, Rosenberg NA. 2008. Mathematical properties of the measure of linkage disequilibrium. Theor Popul Biol 74:130–137.

34. Lin M, Kussell E. 2019. Inferring bacterial recombination rates from large-scale sequencing datasets. Nat Methods 16:199–204.

35. Doroghazi JR, Buckley DH. 2010. Widespread homologous recombination within and between Streptomyces species. ISME J 4:1136–1143.

36. Carja O, Liberman U, Feldman MW. 2014. Evolution in changing environments: Modifiers of mutation, recombination, and migration. Proc Natl Acad Sci 111:17935–17940.

37. Hanage WP. 2016. Not So Simple After All: Bacteria, Their Population Genetics, and Recombination. Cold Spring Harb Perspect Biol 8:a018069.

38. Didelot X, Maiden MCJ. 2010. Impact of recombination on bacterial evolution. Trends Microbiol 18:315–322.

39. Luo H, Gao F, Lin Y. 2015. Evolutionary conservation analysis between the essential and nonessential genes in bacterial genomes. Sci Rep 5:13210.

40. Dilucca M, Cimini G, Giansanti A. 2018. Essentiality, conservation, evolutionary pressure and codon bias in bacterial genomes. Gene 663:178–188.

41. Aguilar-Rodríguez J, Wagner A. 2018. Metabolic Determinants of Enzyme Evolution in a Genome-Scale Bacterial Metabolic Network. Genome Biol Evol 10:3076–3088.

42. Zhong C, Han M, Yu S, Yang P, Li H, Ning K. 2018. Pan-genome analyses of 24 Shewanella strains re-emphasize the diversification of their functions yet evolutionary dynamics of metal-reducing pathway. Biotechnol Biofuels 11:193.

43. Petersen L, Bollback JP, Dimmic M, Hubisz M, Nielsen R. 2007. Genes under positive selection in Escherichia coli. Genome Res 17:1336–1343.

44. Rabby A, Chakraborty S, Rahman A, Shakila Rahman S, Soad S, Fatima Chanda K, Chakravorty R. 2015. Identification of the positively selected genes governing host-pathogen arm race in Vibrio sp. through comparative genomics approach. Biojournal Sci Technol 2.

45. Rodriguez-Valera F, Martin-Cuadrado A-B, Rodriguez-Brito B, Pašić L, Thingstad TF, Rohwer F, Mira A. 2009. Explaining microbial population genomics through phage predation. Nat Rev Microbiol 7:828–836.

46. Cordero OX, Polz MF. 2014. Explaining microbial genomic diversity in light of evolutionary ecology. Nat Rev Microbiol 12:263–273.

47. Bolger AM, Lohse M, Usadel B. 2014. Trimmomatic: A flexible trimmer for Illumina sequence data. Bioinformatics 30:2114–2120.

48. Peng Y, Leung HCM, Yiu SM, Chin FYL. 2012. IDBA-UD: a de novo assembler for single-cell and metagenomic sequencing data with highly uneven depth. Bioinformatics 28:1420–8.

49. Probst AJ, Castelle CJ, Singh A, Brown CT, Anantharaman K, Sharon I, Hug LA, Burstein D, Emerson JB, Thomas BC, Banfield JF. 2017. Genomic resolution of a cold subsurface aquifer community provides metabolic insights for novel microbes adapted to high CO 2 concentrations. Environ Microbiol 19:459–474.

50. Kang DD, Froula J, Egan R, Wang Z. 2015. MetaBAT, an efficient tool for accurately reconstructing single genomes from complex microbial communities. PeerJ 3:e1165.

51. Wu Y-W, Simmons BA, Singer SW. 2016. MaxBin 2.0: an automated binning algorithm to recover genomes from multiple metagenomic datasets. Bioinformatics 32:605–607.

52. Alneberg J, Bjarnason BS, de Bruijn I, Schirmer M, Quick J, Ijaz UZ, Lahti L, Loman NJ, Andersson AF, Quince C. 2014. Binning metagenomic contigs by coverage and composition. Nat Methods 11:1144–1146.

53. Lin H-H, Liao Y-C. 2016. Accurate binning of metagenomic contigs via automated clustering sequences using information of genomic signatures and marker genes. Sci Rep 6:24175.

54. Graham ED, Heidelberg JF, Tully BJ. 2017. BinSanity: unsupervised clustering of environmental microbial assemblies using coverage and affinity propagation. PeerJ 5:e3035.

55. Lu YY, Chen T, Fuhrman JA, Sun F. 2016. COCACOLA: binning metagenomic contigs using sequence COmposition, read CoverAge, CO-alignment and paired-end read LinkAge. Bioinformatics btw290.

56. Song WZ, Thomas T. 2017. Binning-refiner: Improving genome bins through the combination of different binning programs. Bioinformatics 33:1873–1875.

57. Uritskiy G V., DiRuggiero J, Taylor J. 2018. MetaWRAP—a flexible pipeline for genome-resolved metagenomic data analysis. Microbiome 6:158.

58. Sieber CMK, Probst AJ, Sharrar A, Thomas BC, Hess M, Tringe SG, Banfield JF. 2018. Recovery of genomes from metagenomes via a dereplication, aggregation and scoring strategy. Nat Microbiol 3:836–843.

59. Olm MR, Brown CT, Brooks B, Banfield JF. 2017. DRep: A tool for fast and accurate genomic comparisons that enables improved genome recovery from metagenomes through de-replication. ISME J 11:2864–2868.

60. Karst SM, Kirkegaard RH, Albertsen M. 2016. Mmgenome: a Toolbox for Reproducible Genome Extraction From Metagenomes. bioRxiv 059121.

61. Parks DH, Rinke C, Chuvochina M, Chaumeil PA, Woodcroft BJ, Evans PN, Hugenholtz P, Tyson GW. 2017. Recovery of nearly 8,000 metagenome-assembled genomes substantially expands the tree of life. Nat Microbiol 2:1533–1542.

62. Bosi E, Donati B, Galardini M, Brunetti S, Sagot MF, Lió P, Crescenzi P, Fani R, Fondi M. 2015. MeDuSa: A multi-draft based scaffolder. Bioinformatics 31:2443–2451.

63. Parks DH, Imelfort M, Skennerton CT, Hugenholtz P, Tyson GW. 2015. CheckM: assessing the quality of microbial genomes recovered from isolates, single cells, and metagenomes. Genome Res 25:1043–55.

64. Nayfach S, Rodriguez-Mueller B, Garud N, Pollard KS. 2016. An integrated metagenomics pipeline for strain profiling reveals novel patterns of bacterial transmission and biogeography. Genome Res 26:1612–1625.

65. Chaumeil P-A, Mussig AJ, Hugenholtz P, Parks DH. 2019. GTDB-Tk: a toolkit to classify genomes with the Genome Taxonomy Database. Bioinformatics https://doi.org/10.1093/bioinformatics/btz848.

66. Hyatt D, Chen G-L, Locascio PF, Land ML, Larimer FW, Hauser LJ. 2010. Prodigal: prokaryotic gene recognition and translation initiation site identification. BMC Bioinformatics 11:119.

67. Altschul SF, Gish W, Miller W, Myers EW, Lipman DJ. 1990. Basic local alignment search tool. J Mol Biol 215:403–410.

68. Sayers EW, Cavanaugh M, Clark K, Ostell J, Pruitt KD, Karsch-Mizrachi I. 2019. GenBank. Nucleic Acids Res 47:D94–D99.

69. 2019. UniProt: a worldwide hub of protein knowledge. Nucleic Acids Res 47:D506–D515.

70. Ogata H, Goto S, Sato K, Fujibuchi W, Bono H, Kanehisa M. 1999. KEGG: Kyoto encyclopedia of genes and genomes. Nucleic Acids Res 27:29–34.

71. Punta M, Coggill PC, Eberhardt RY, Mistry J, Tate J, Boursnell C, Pang N, Forslund K, Ceric G, Clements J, Heger A, Holm L, Sonnhammer ELL, Eddy SR, Bateman A, Finn RD. 2012. The Pfam protein families database. Nucleic Acids Res 40:D290–D301.

72. Huerta-Cepas J, Szklarczyk D, Forslund K, Cook H, Heller D, Walter MC, Rattei T, Mende DR, Sunagawa S, Kuhn M, Jensen LJ, von Mering C, Bork P. 2016. eggNOG 4.5: a hierarchical orthology framework with improved functional annotations for eukaryotic, prokaryotic and viral sequences. Nucleic Acids Res 44:D286–D293.

73. Doron S, Melamed S, Ofir G, Leavitt A, Lopatina A, Keren M, Amitai G, Sorek R. 2018. Systematic discovery of antiphage defense systems in the microbial pangenome. Science (80-) 359:eaar4120.

74. Parks DH, Chuvochina M, Waite DW, Rinke C, Skarshewski A, Chaumeil P-A, Hugenholtz P. 2018. A standardized bacterial taxonomy based on genome phylogeny substantially revises the tree of life. Nat Biotechnol https://doi.org/10.1038/nbt.4229.

75. Stamatakis A. 2014. RAxML version 8: a tool for phylogenetic analysis and post-analysis of large phylogenies. Bioinformatics 30:1312–1313.

76. Darriba D, Taboada GL, Doallo R, Posada D. 2011. ProtTest 3: fast selection of best-fit models of protein evolution. Bioinformatics 27:1164–1165.

77. Letunic I, Bork P. 2016. Interactive tree of life (iTOL) v3: an online tool for the display and annotation of phylogenetic and other trees. Nucleic Acids Res 44:W242–W245.

78. Li H, Durbin R. 2010. Fast and accurate long-read alignment with Burrows-Wheeler transform. Bioinformatics 26:589–95.

79. Li H, Handsaker B, Wysoker A, Fennell T, Ruan J, Homer N, Marth G, Abecasis G, Durbin R. 2009. The Sequence Alignment/Map format and SAMtools. Bioinformatics 25:2078–2079.

80. Team RC. 2014. R: A language and environment for statistical computing. R Foundation for Statistical Computing, Vienna, Austria.

81. Gloor GB, Macklaim JM, Pawlowsky-Glahn V, Egozcue JJ. 2017. Microbiome Datasets Are Compositional: And This Is Not Optional. Front Microbiol 8.

82. Palarea-Albaladejo J, Martín-Fernández JA. 2015. zCompositions — R package for multivariate imputation of left-censored data under a compositional approach. Chemom Intell Lab Syst 143:85–96.

83. Quinn TP, Richardson MF, Lovell D, Crowley TM. 2017. propr: An R-package for Identifying Proportionally Abundant Features Using Compositional Data Analysis. Sci Rep 7:16252.

84. Wei T, Simko V. 2017. R Package “corrplot”: visualization of a correlation matrix.

85. Gabor C, Tamas N. 2006. The igraph software package for complex network research. InterJournal Complex Syst 1695.

86. Oksanen J, Blanchet FG, Friendly M, Kindt R, Legendre P, Mcglinn D, Minchin PR, O’hara RB, Simpson GL, Solymos P, Henry M, Stevens H, Szoecs E, Maintainer HW. 2019. Package “vegan” Title Community Ecology Package. Community Ecol Packag 2:1–297.

87. Nakazawa M. 2021. Fmsb: Functions for Medical Statistics Book with Some Demographic Data. R Package Version 0.7.1. https://cran.r-project.org/package=fmsb.

88. Beck MW. 2017. vif_fun.r. https://gist.github.com/fawda123/4717702.

89. Hamilton NE, Ferry M. 2018. ggtern : Ternary Diagrams Using ggplot2. J Stat Softw 87.

90. Bower CE, Holm-Hansen T. 1980. A Salicylate–Hypochlorite Method for Determining Ammonia in Seawater. Can J Fish Aquat Sci 37:794–798.

91. Grasshoff K, Ehrhardt M, Kremling K. 1983. Methods of Seawater Analysis, 2nd ed. Verlag Chemie, Basel, CH.

92. Eaton A, Clesceri L, Greenberg A, Franson M, American Public Health Association, American Water Works Association, Water Environment Federation. 1998. Standard methods for the examination of water and wastewater, 20th ed. American Public Health Association, Washington D.C.

