## Supplementary Information for "Evolutionary ecology of natural comammox *Nitrospira* populations"

Alejandro Palomo[1], Arnaud Dechesne[1], Otto X. Cordero[2] and Barth F. Smets[1]
[1]Department of Environmental Engineering, Technical University of Denmark, Kgs Lyngby, Denmark
[2]Ralph M. Parsons Laboratory for Environmental Science and Engineering, Department of Civil and Environmental Engineering, Massachusetts Institute of Technology, Cambridge, MA, USA

**Supplementary Information includes:**
Fig. S1 to S13

**Other supplementary materials for this manuscript include the following:**
Table S1 to S10

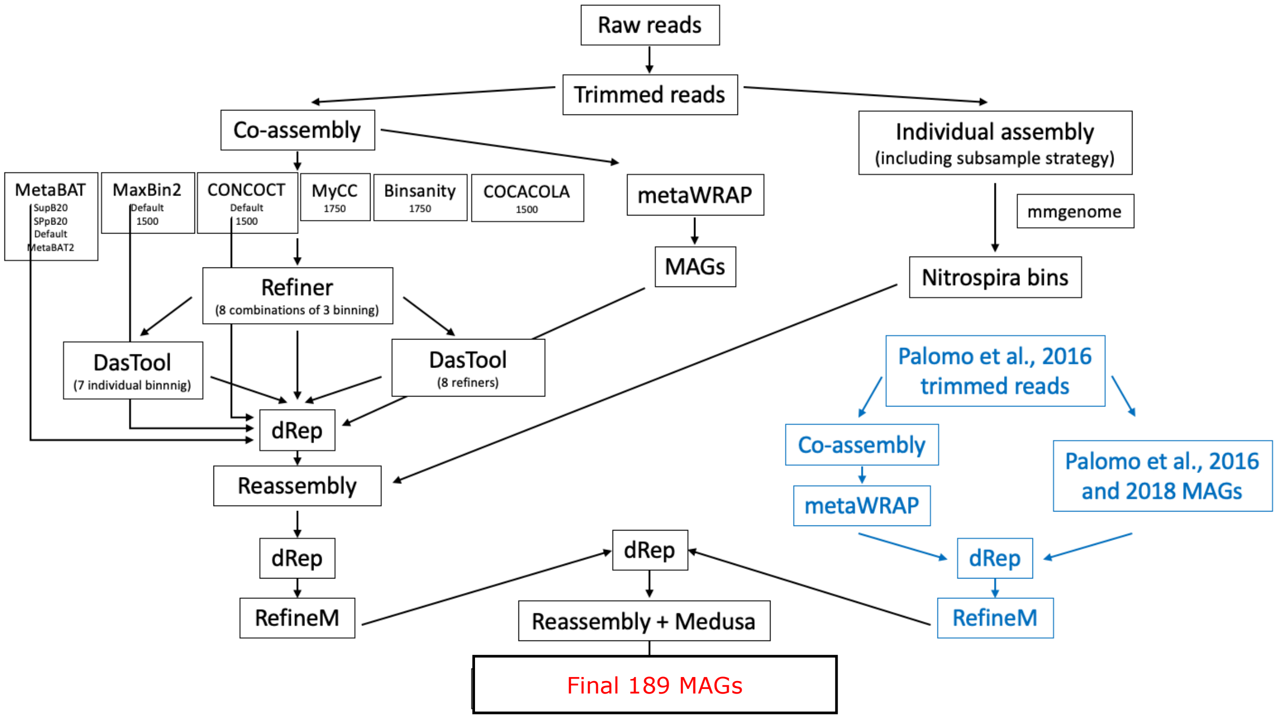


Fig. S1. The implemented workflow for the MAGs recovery from 12 Danish groundwater-fed rapid sand filters. The final genome quality improvement performed with MeDuSa was only applied on the18 *Nitrospira* MAGs.

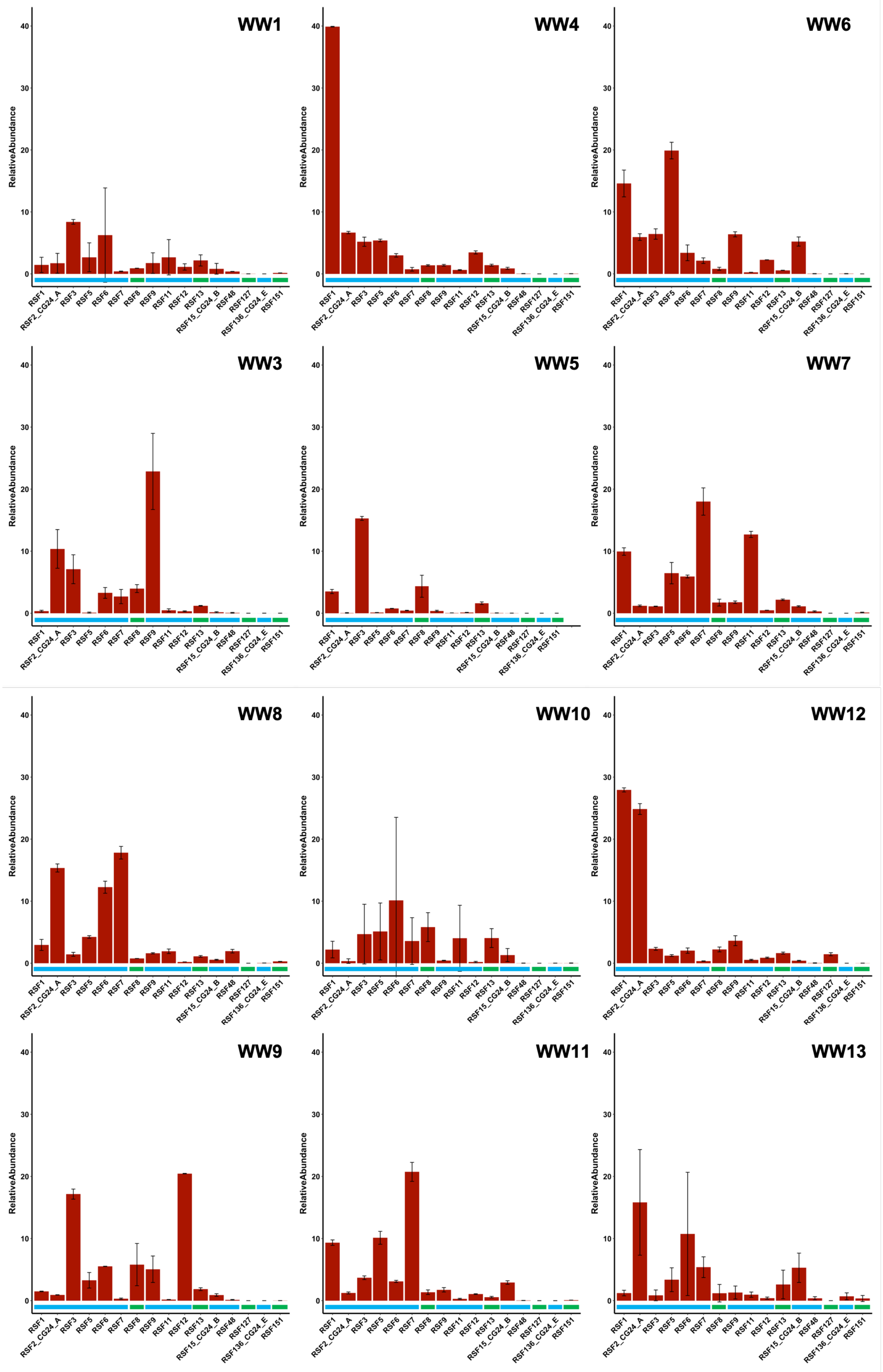


Fig. S2. Average relative abundance (n=2) of 16 *Nitrospira* species in 12 waterworks. Comammox and canonical *Nitrospira* species are denoted in blue and green, respectively.

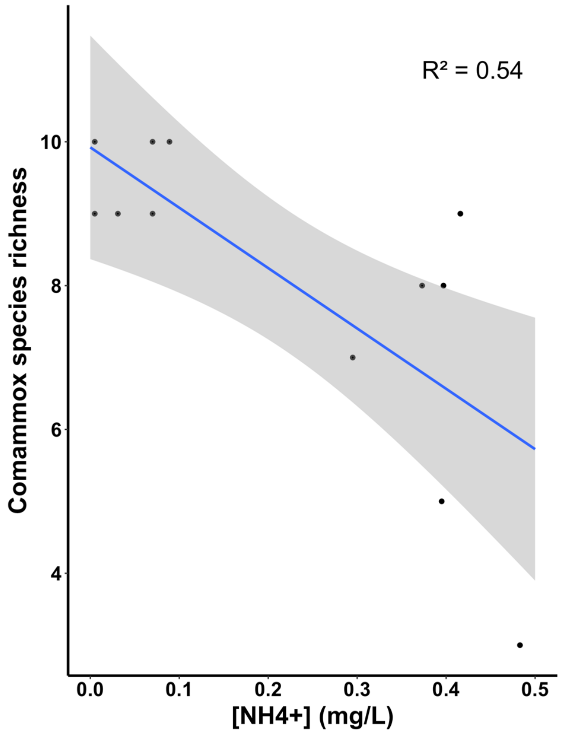


Fig. S3. Relationship between the ammonium concentration and the comammox species richness in each waterworks (number of species with abundances > 0.5%). Blue line shows the linear regression with shadowed region indicating 95% confidence intervals for the slope (p value for $R^2 < 0.01$).

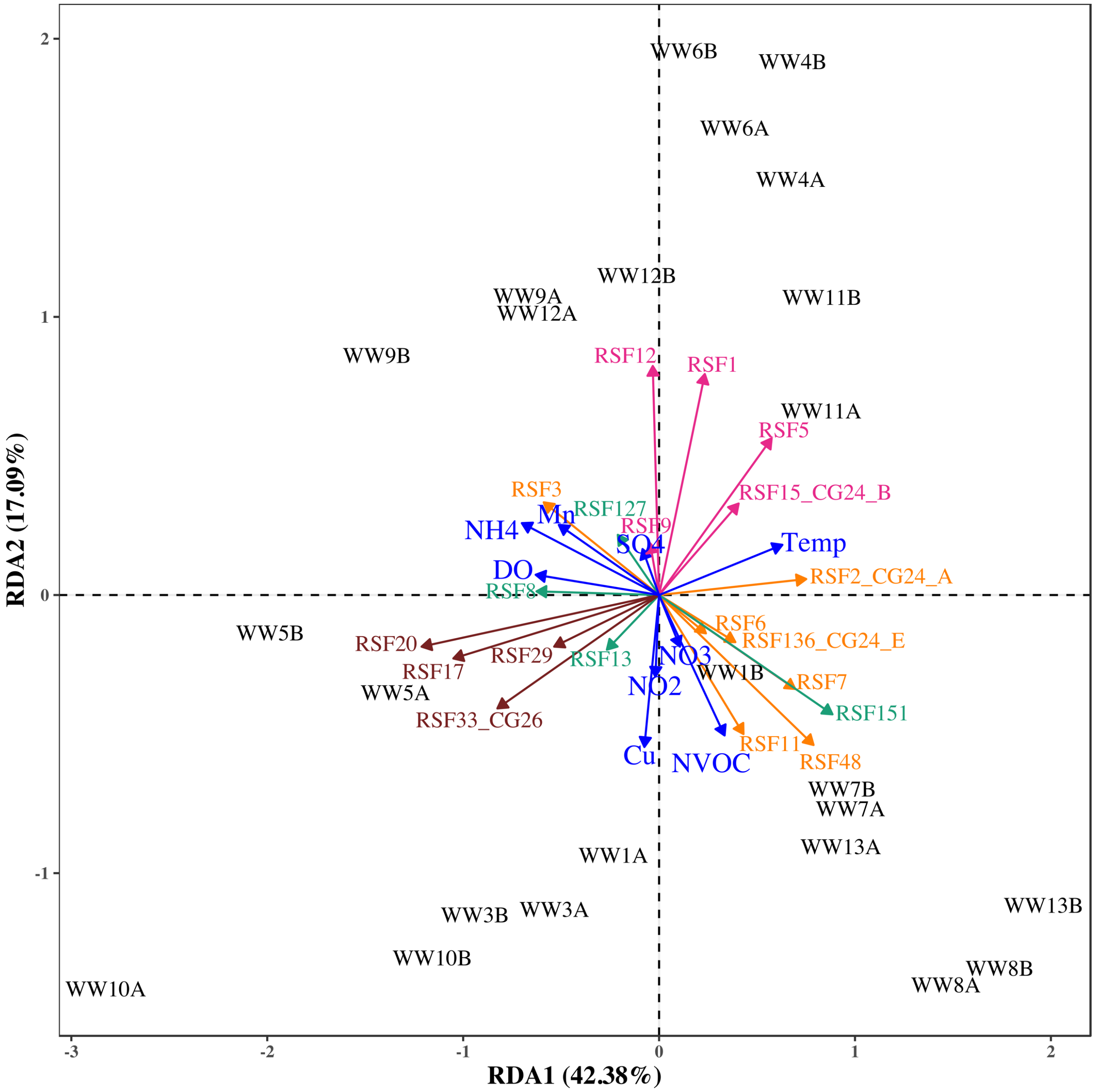


Fig. S4. Redundancy analysis (RDAof the relationship between water chemistry and the species abundance in 12 waterworks. Arrows show the association of individual species (different colours) and water chemistry variables (blue arrows) with each axis. Species colour code: pink: comammox clade A; orange: comammox clade B; green: canonical *Nitrospira*; brown: ßcanonical AOB.

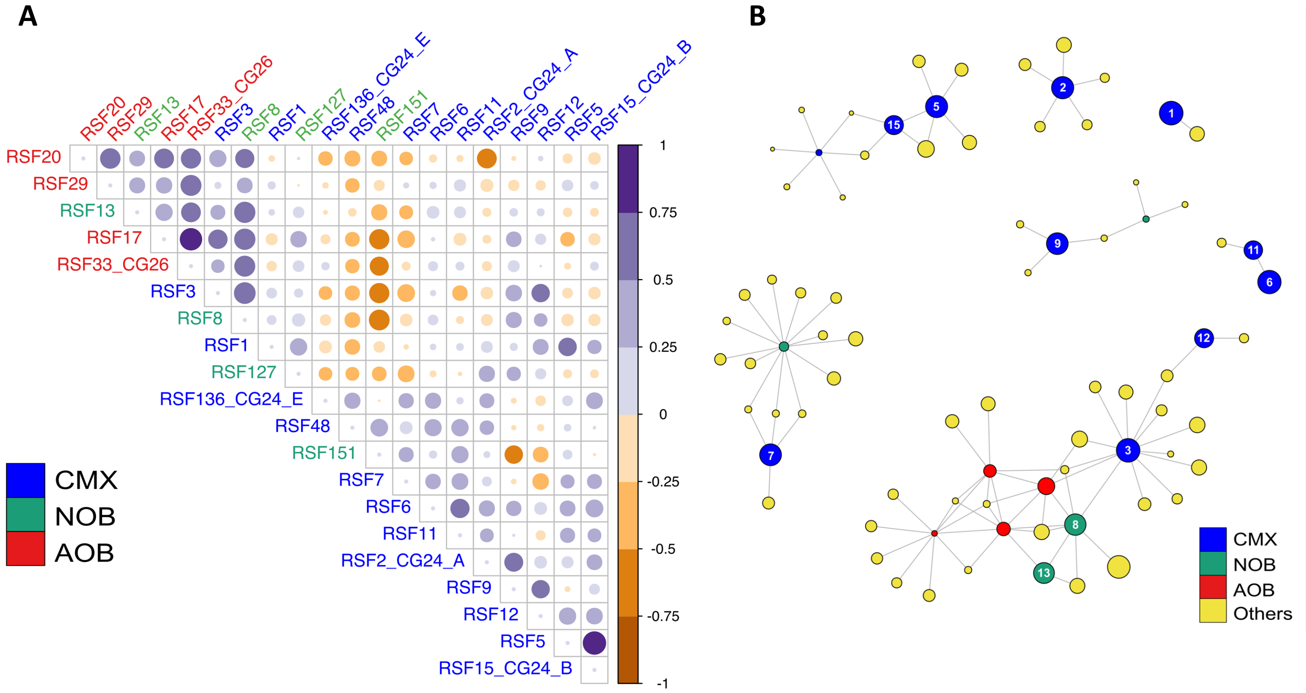


Fig. S5. A) Correlogram showing proportionality for centred log-transformed abundances of the *Nitrospira* and AOB species across the 12 waterworks. Colour indicates whether the correlation is positive (purple) or negative (brown). Size and darkness of the circles indicate the strength of the proportionality, with stronger proportionality being larger and darker than weaker ones. B) Network analysis revealing the co-occurrence patterns among species present in the studied waterworks. Each node represents a species, and are coloured according to species type (lineage II canonical *Nitrospira*, green; comammox, blue; AOB, red; other bacteria, yellow). A connection represents a strong proportionality ($\rho$>0.55 and FDR < 0.05). The size of each node is proportional to the average species log-transformed abundance. Only nodes connected with *Nitrospira* species are shown. The MAG identity of the *Nitrospira* species is displayed with a number on top of the node (e.g.: 1 stands for the species RSF1).

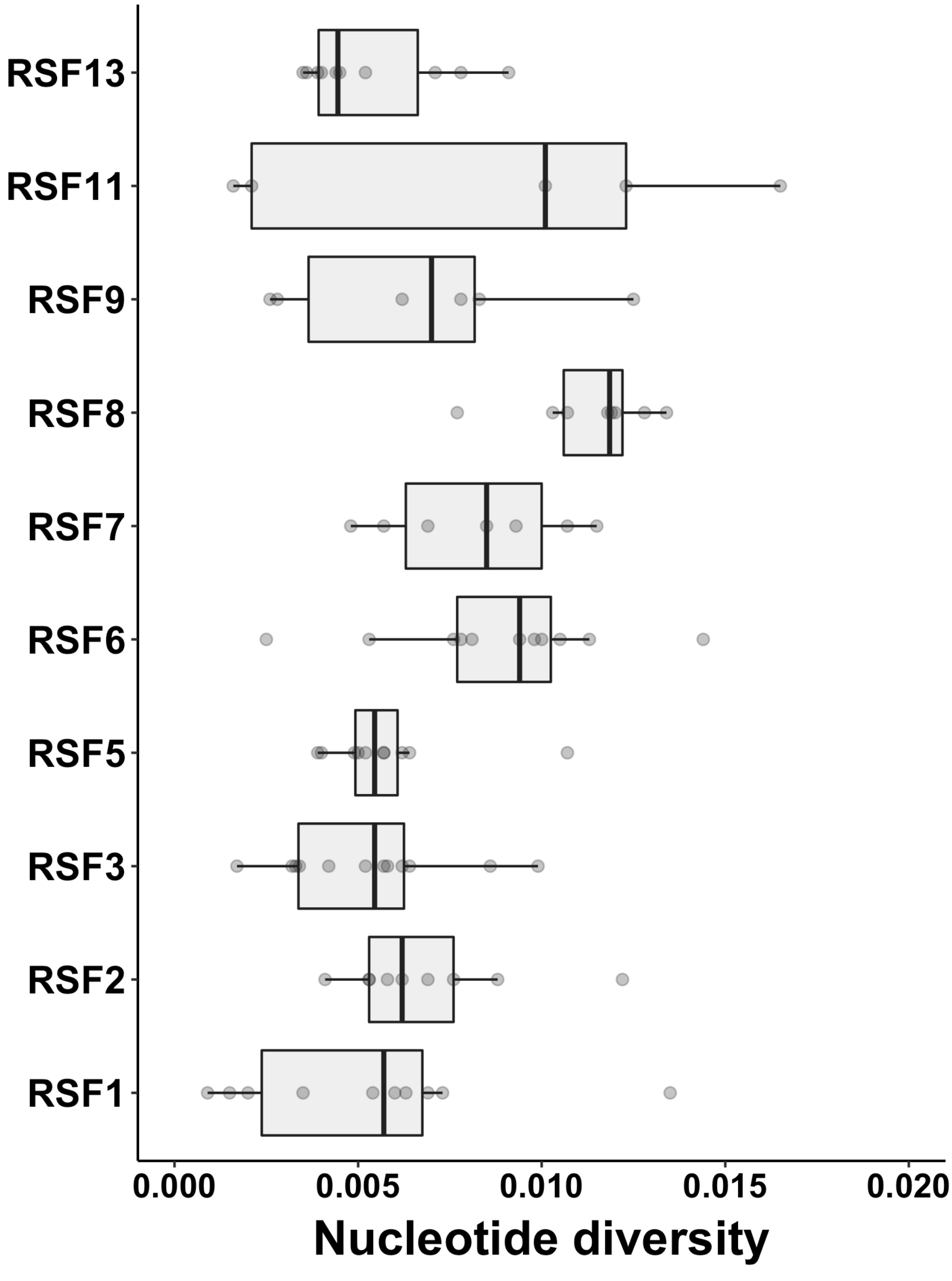


Fig. S6. Boxplot showing the nucleotide diversity values for each *Nitrospira* species in the studied waterworks. Only species with more than two data points are shown.

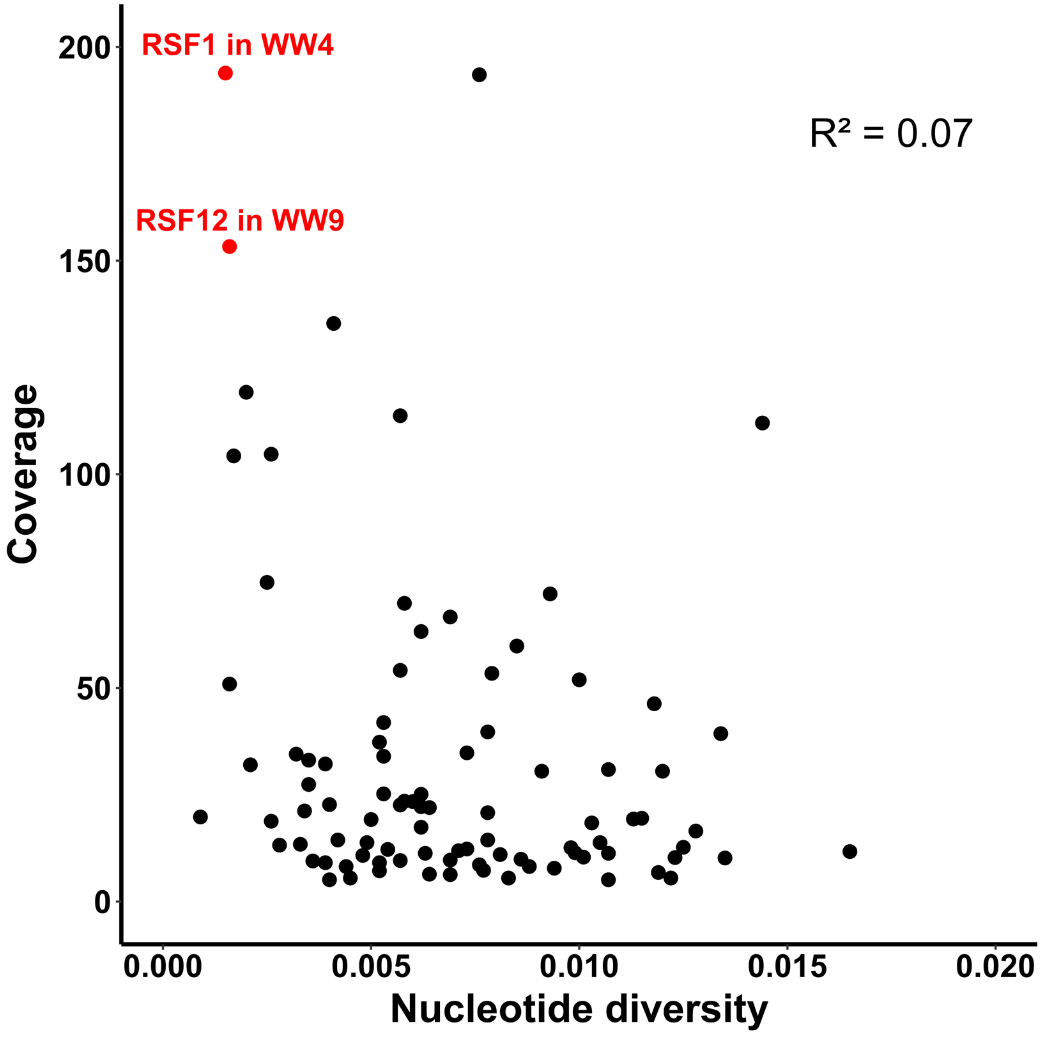


Fig. S7. The relationship between nucleotide diversity and coverage for each *Nitrospira* population in each waterworks. The mean nucleotide diversity and the mean coverage across species in each waterworks is shown. Local populations with high coverage and low nucleotide diversity are coloured in red. (linear regression; R2 = 0.07; p > 0.01).

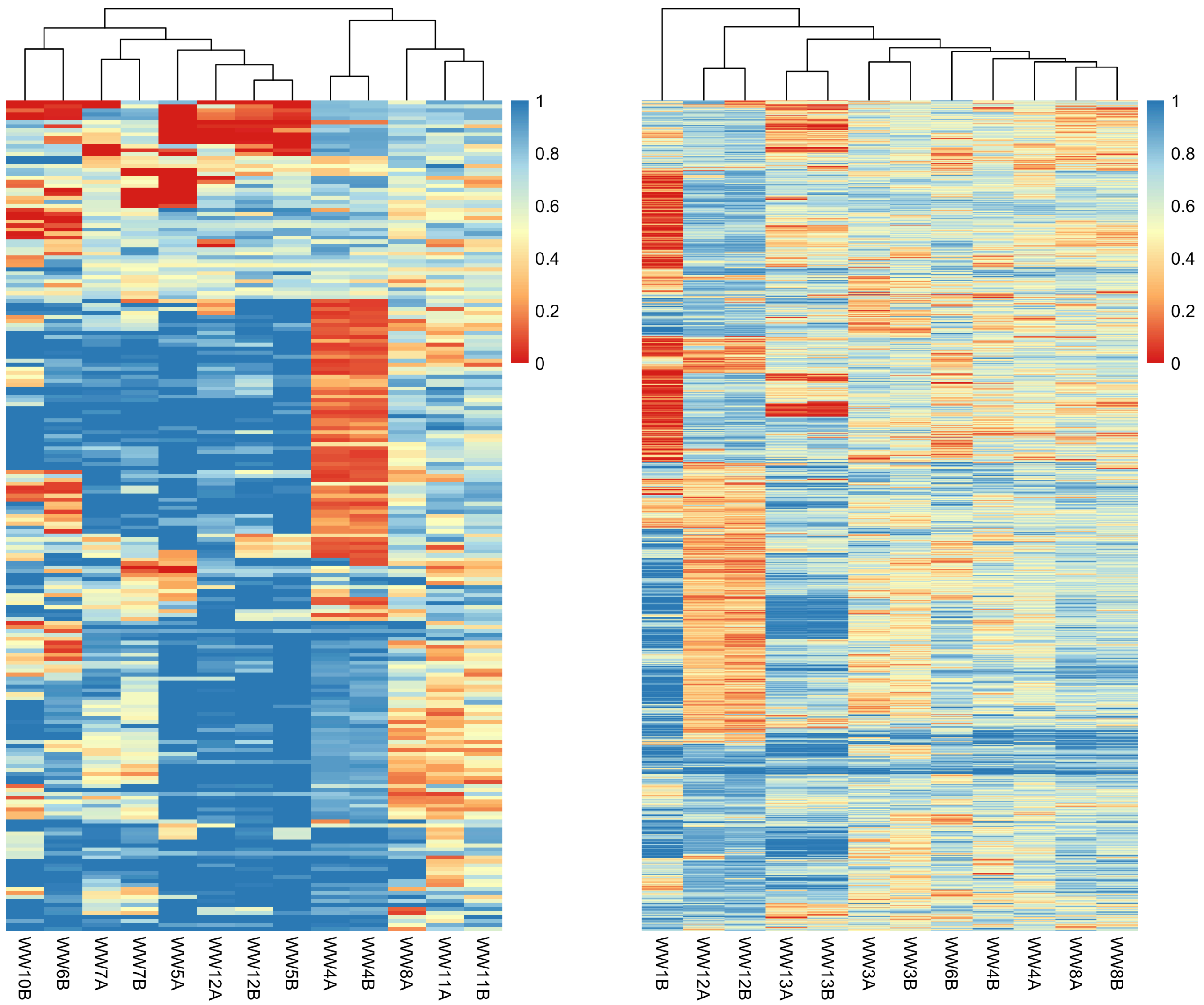
WW10B
WW6B
WW7A
WW7B
WW5A
WW12A
WW12B
WW5B
WW4A
WW4B
WW8A
WW11A
WW11B
1
0.8
0.6
0.4
0.2
0
WW1B
WW12A
WW12B
WW13A
WW13B
WW3A
WW3B
WW6B
WW4B
WW4A
WW8A
WW8B
1
0.8
0.6
0.4
0.2
0

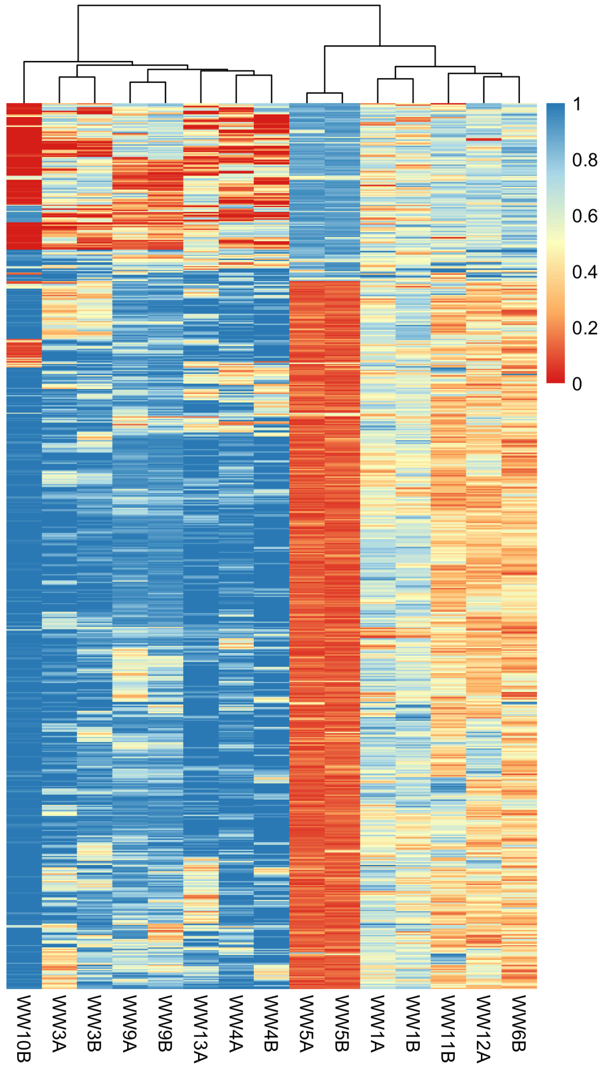
WW10B
WW3A
WW3B
WW9A
WW9B
WW13A
WW4A
WW4B
WW5A
WW5B
WW1A
WW1B
WW11B
WW12A
WW6B
1
0.8
0.6
0.4
0.2
0

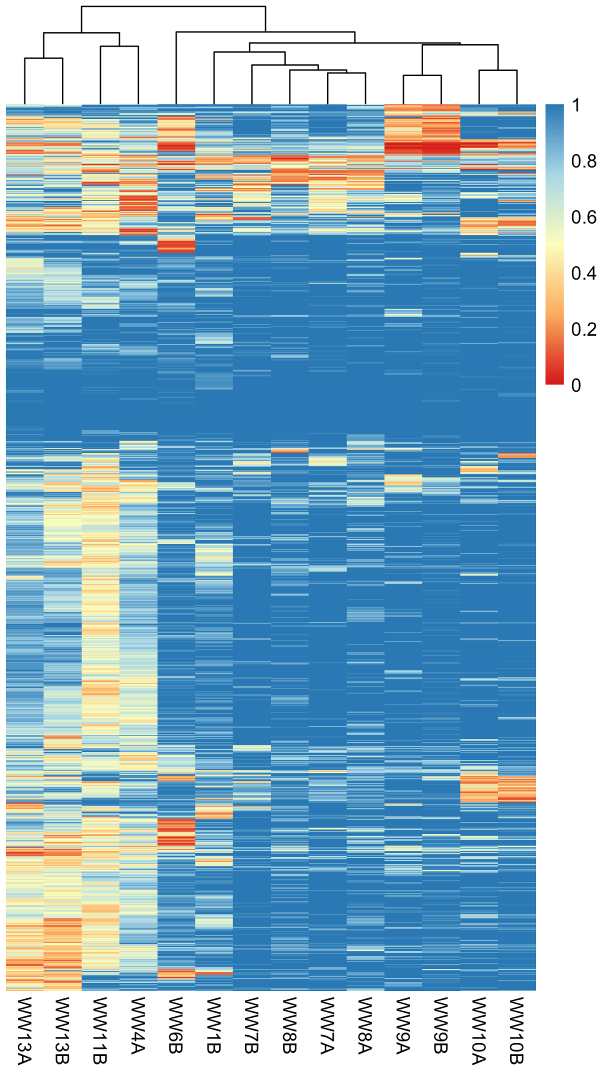
WW13A
WW13B
WW11B
WW4A
WW6B
WW1B
WW7B
WW8B
WW7A
WW8A
WW9A
WW9B
WW10A
WW10B
1
0.8
0.6
0.4
0.2
0

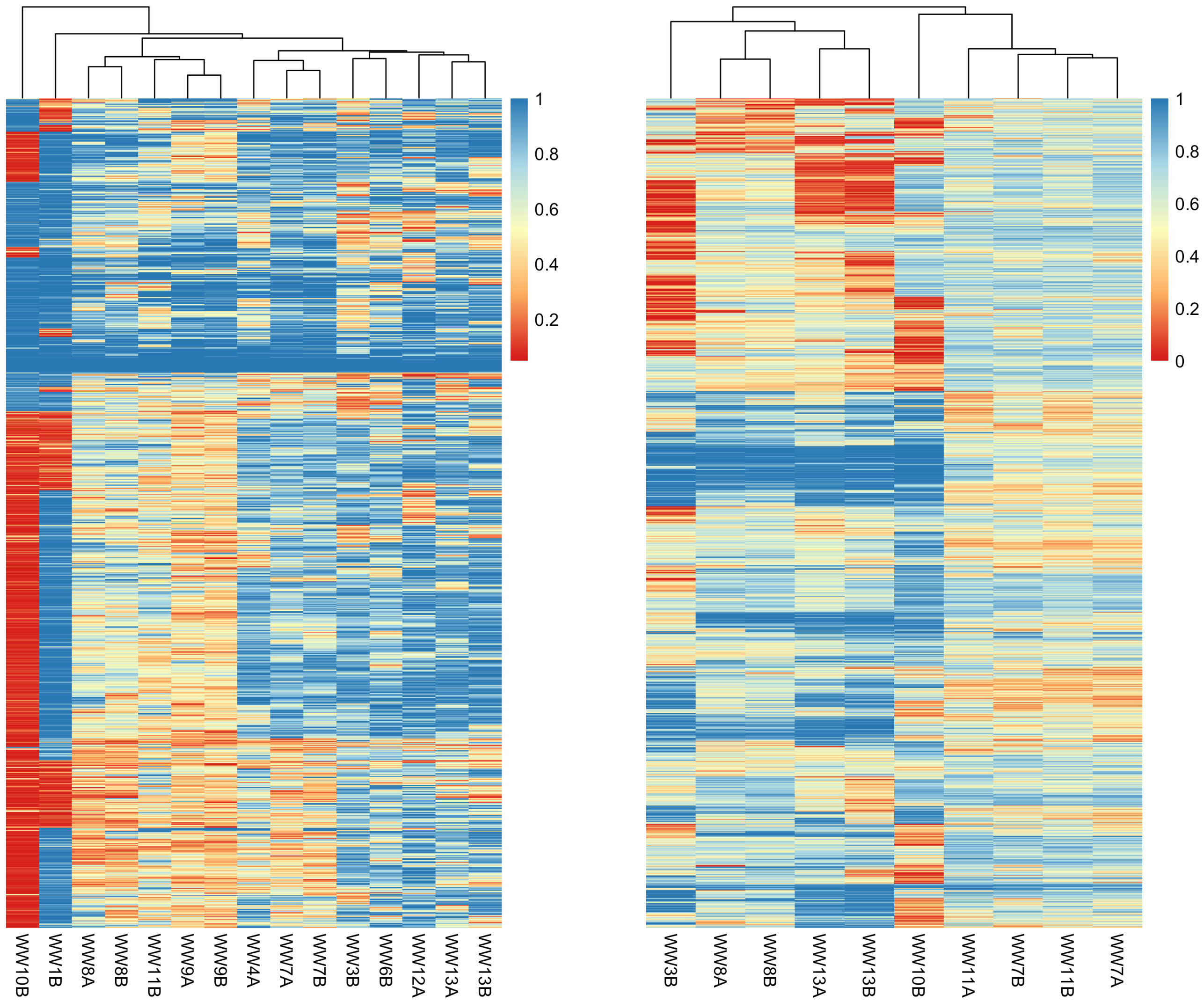

1
0.8
0.6
0.4
0.2
WW10B
WW1B
WW8A
WW8B
WW11B
WW9A
WW9B
WW4A
WW7A
WW7B
WW3B
WW6B
WW12A
WW13A
WW13B
1
0.8
0.6
0.4
0.2
0
WW3B
WW8A
WW8B
WW13A
WW13B
WW10B
WW11A
WW7B
WW11B
WW7A

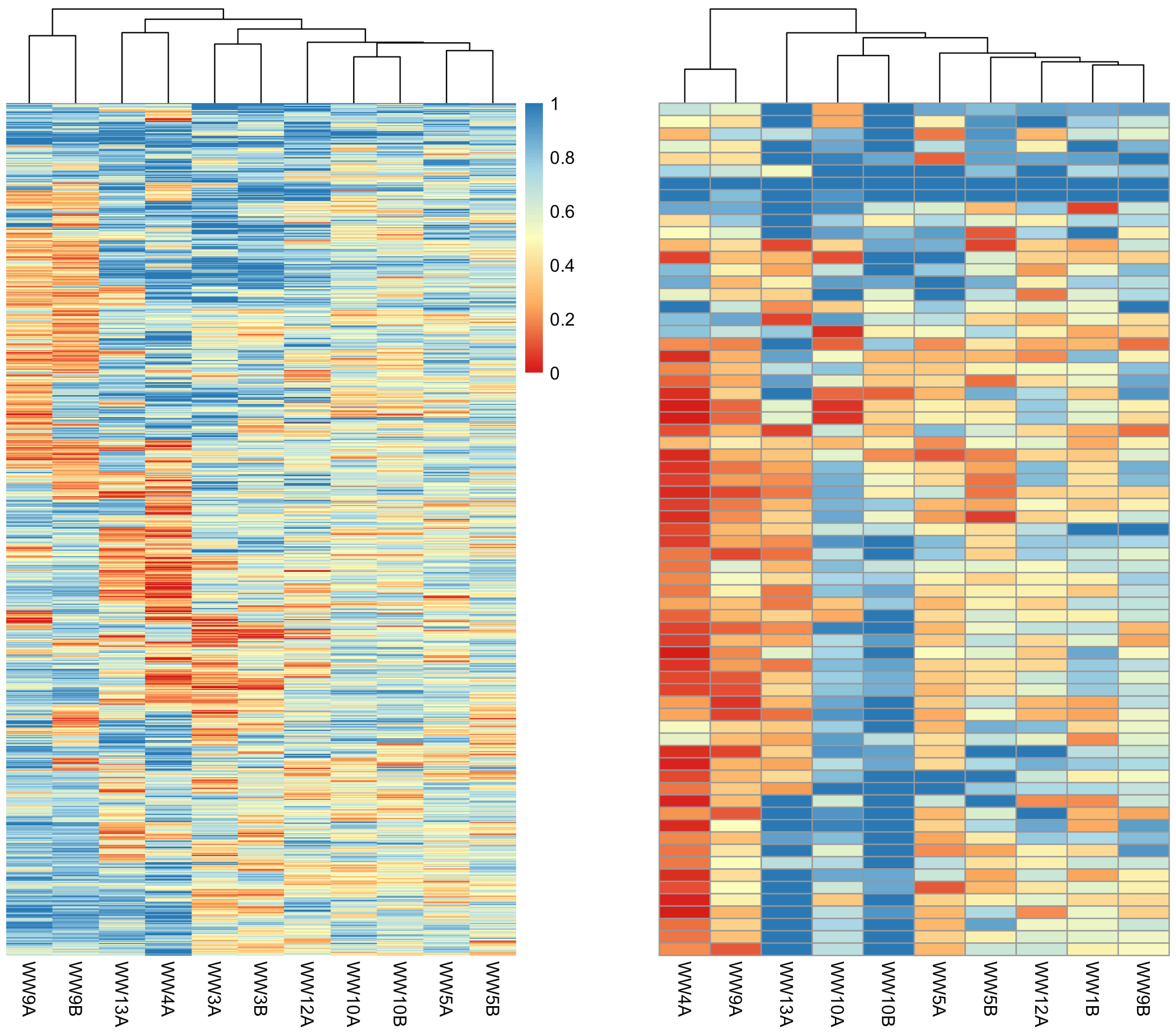


Fig. S8. Heatmap of the major allele frequency for common SNPs (rows) in the waterworks populations (columns; only included the populations with coverage > 5) of the different *Nitrospira* species. Colour represents the frequency of major allele of each SNP from 0 (red) to 1 (blue). Dendrograms group populations based on the frequency similarity. The order of the eights plots from left to right and top to bottom corresponds to the following species: RSF1, RSF2, RSF3, RSF5, RSF6, RSF7, RSF8, RSF13 (this analysis only includes the RSF species with a coverage > 5 in at least 10 samples).

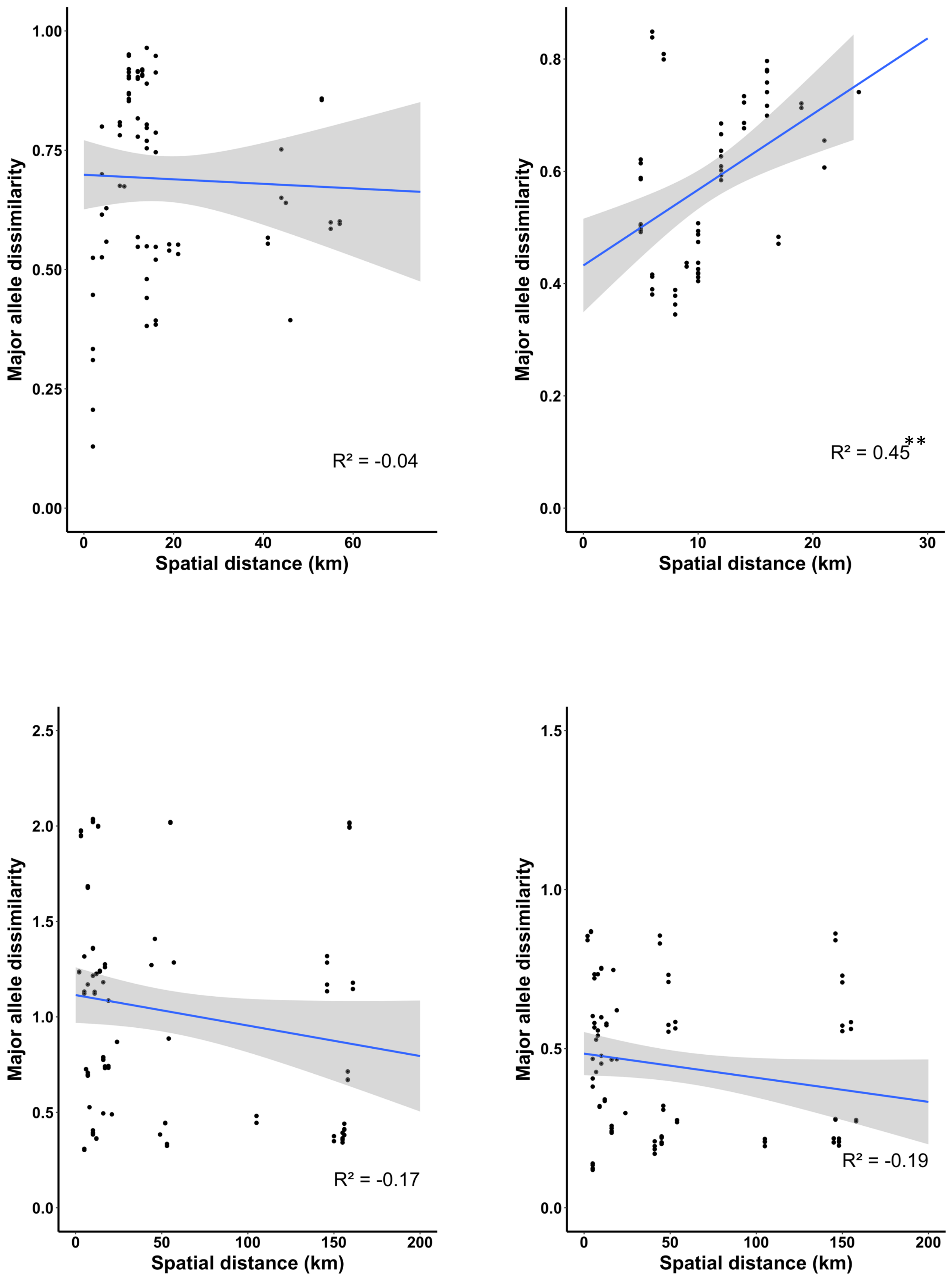

Major allele dissimilarity
Spatial distance (km)
R² = -0.04
Major allele dissimilarity
Spatial distance (km)
R² = 0.45**
Major allele dissimilarity
Spatial distance (km)
R² = -0.17
Major allele dissimilarity
Spatial distance (km)
R² = -0.19

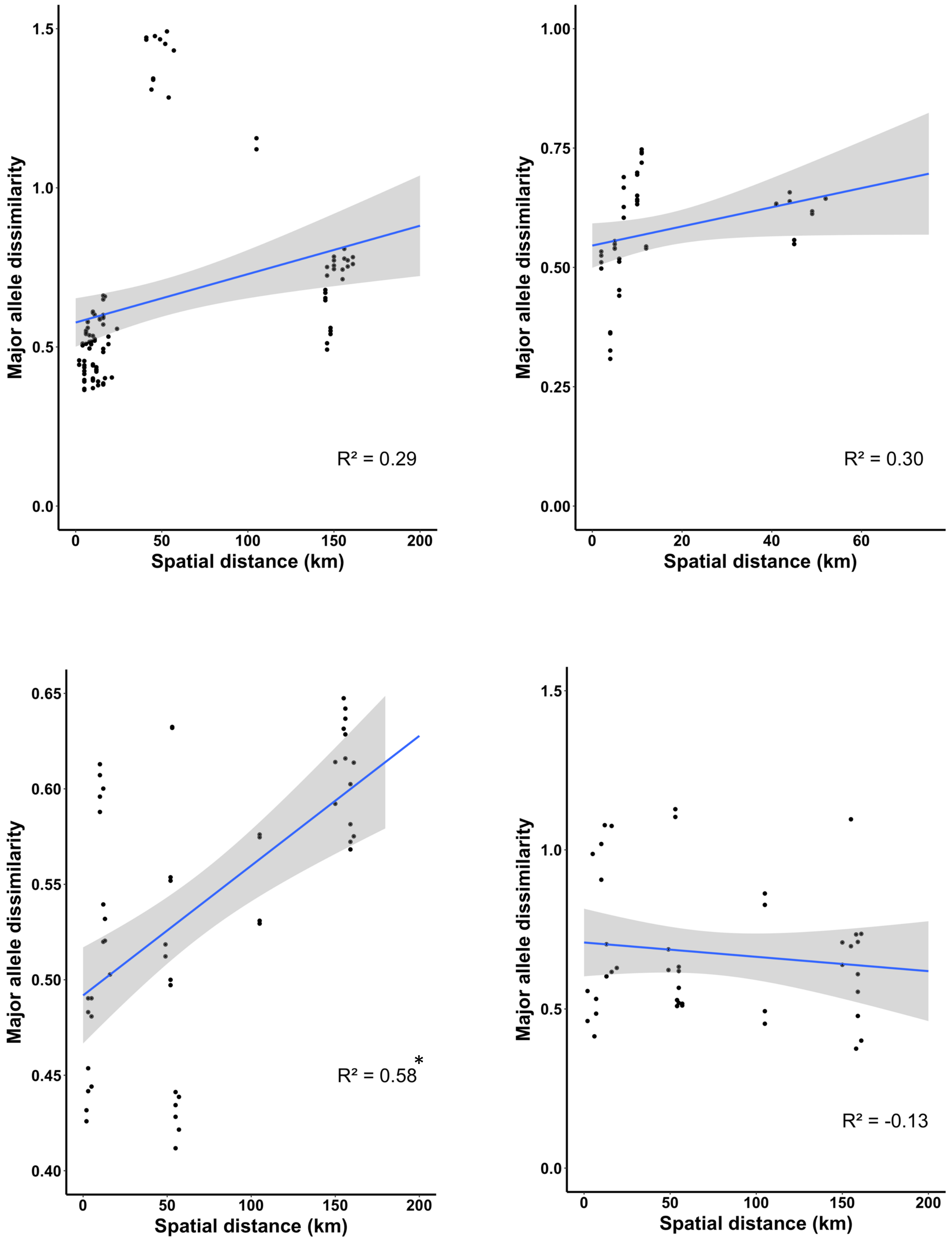
Major allele dissimilarity
Spatial distance (km)
R² = 0.29
R² = 0.30
R² = 0.58*
R² = -0.13

Fig. S9. Relationship between waterworks dissimilarity based on major allele frequency of common SNPs and the geographic distance of the waterworks. The dissimilarities between populations in pairs of waterworks are calculated using the Jaccard index from a matrix of major allele frequencies for common SPNs across the waterworks: the value 0 means that the two population in the two waterworks have the same allele profile. The Mantel test was used to test the strength and significance of correlations (R2 denotes the Mantel statistic *r;* * denotes p < 0.05 and ** denotes p < 0.01). Blue line shows the linear regression with shadowed region indicating 95% confidence intervals for the slope. The order of the plots from left to right and top to bottom corresponds to: RSF1, RSF2, RSF3, RSF5, RSF6, RSF7, RSF8, RSF13.

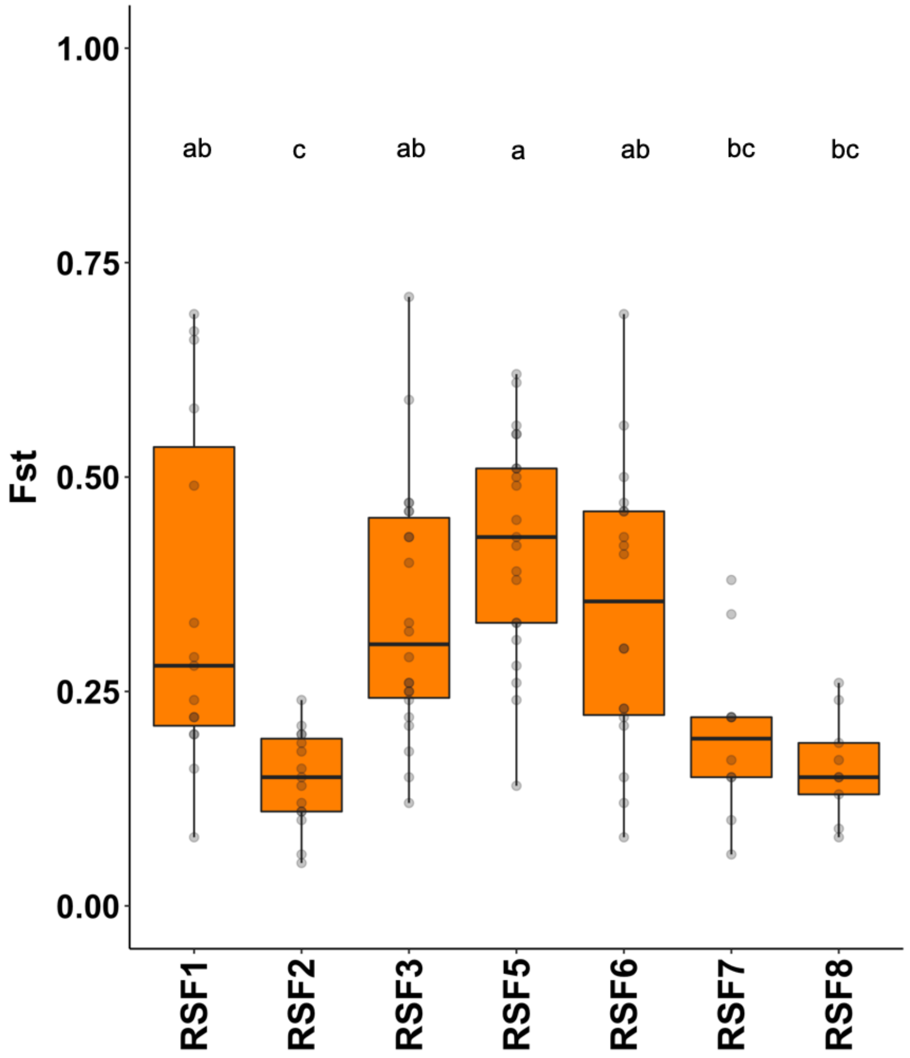


Fig. S10. Boxplot showing the $F_{ST}$ values for each species (each dot represents the $F_{ST}$ value measured as the differences in allele frequencies between populations of the same species found in two distinct waterworks). Only species with more than two data points are shown. Differences between the mean $F_{ST}$ were assessed by a Dunn's test; same letter have means not significantly different from each other ($p < 0.01$).

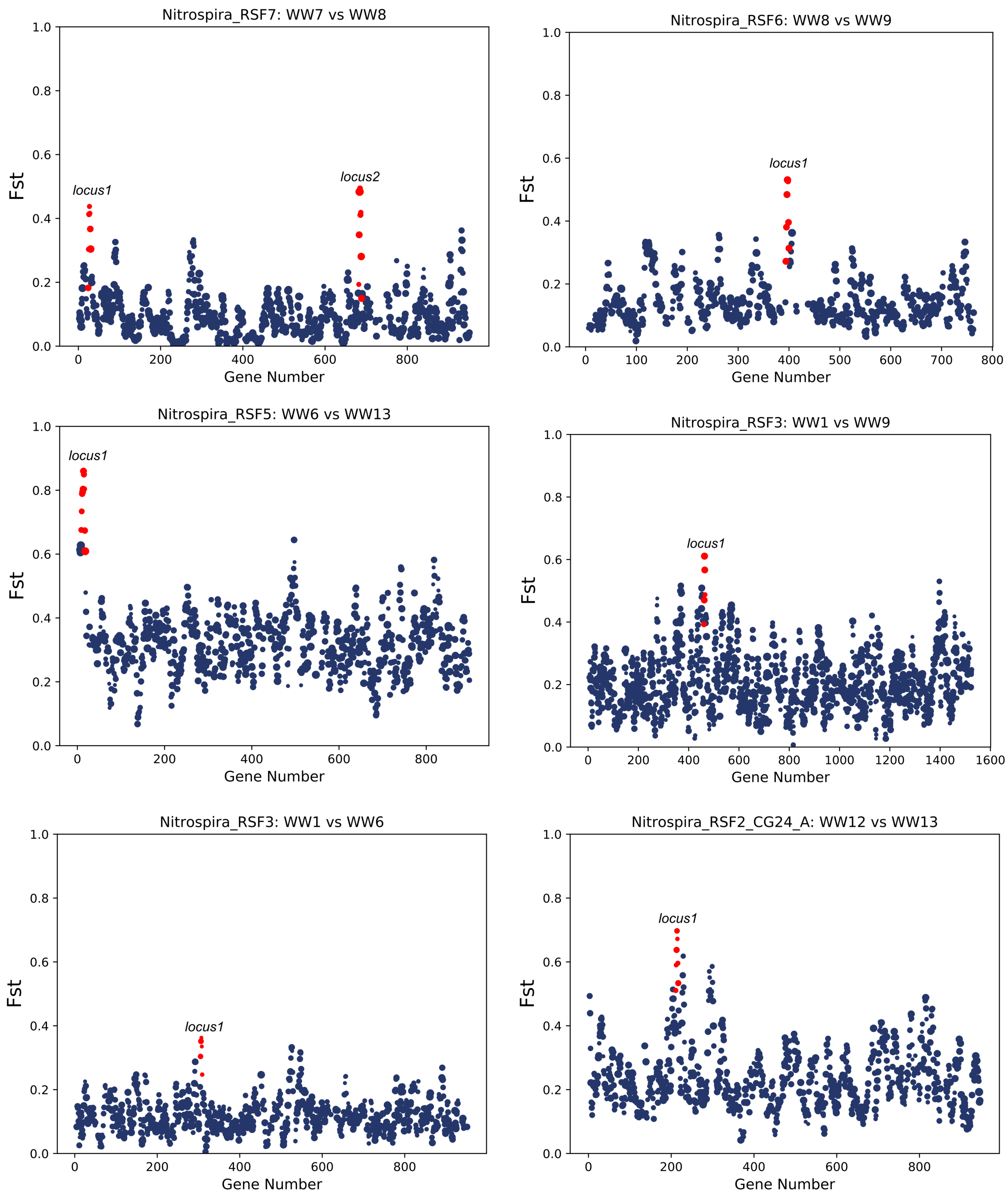

Nitrospira_RSF7: WW7 vs WW8
Fst
Gene Number
locus1
locus2
Nitrospira_RSF6: WW8 vs WW9
Fst
Gene Number
locus1
Nitrospira_RSF5: WW6 vs WW13
Fst
Gene Number
locus1
Nitrospira_RSF3: WW1 vs WW9
Fst
Gene Number
locus1
Nitrospira_RSF3: WW1 vs WW6
Fst
Gene Number
locus1
Nitrospira_RSF2_CG24_A: WW12 vs WW13
Fst
Gene Number
locus1

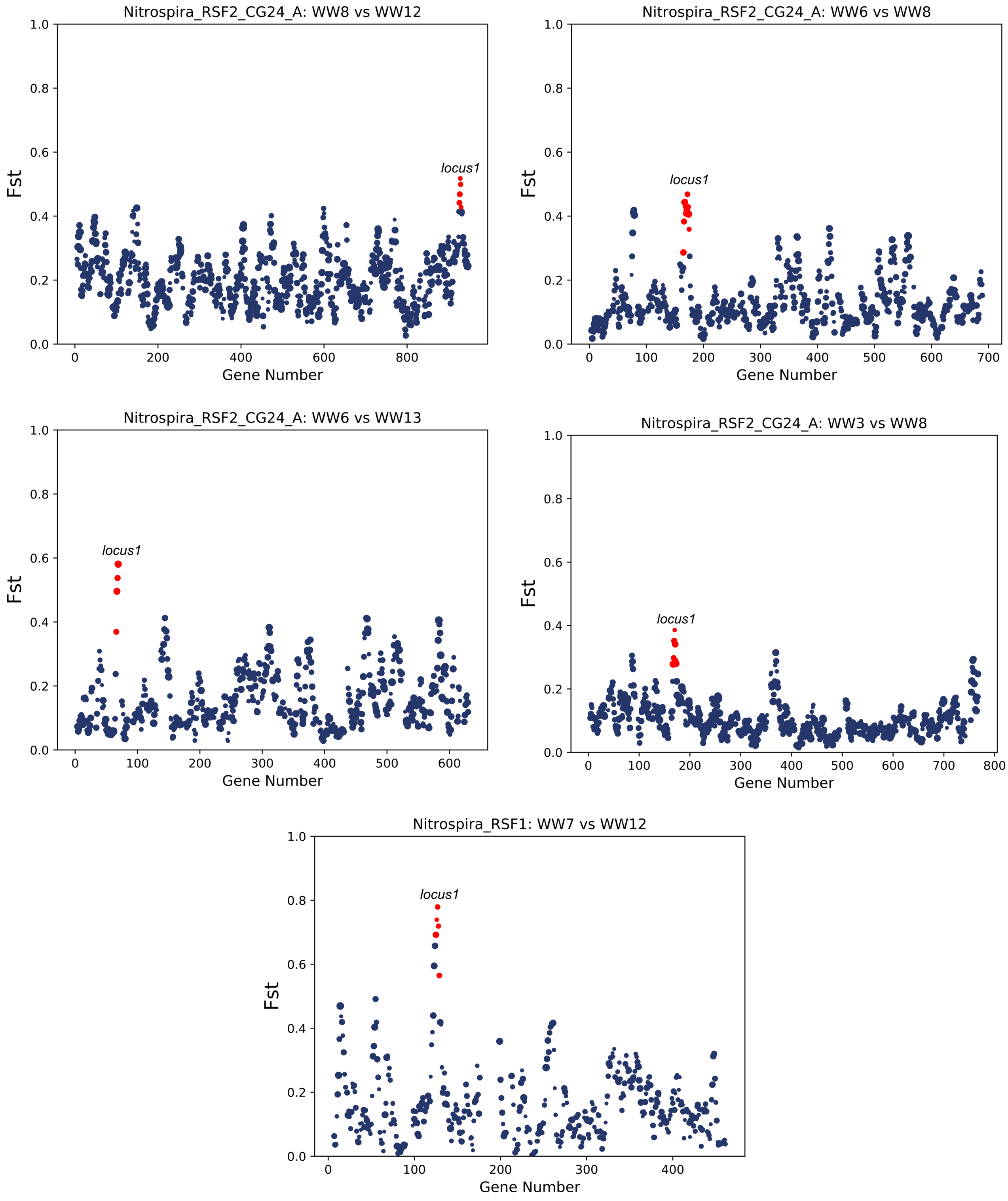


Fig. S10. Highly different genomic loci between waterworks. Values of $F_{ST}$ for genes across the genomes of the bacterial populations. Each point is a gene, and the size of the point is determined by the number of SNPs within that gene. Loci with significantly higher $F_{ST}$ than the background are highlighted in red. (Here are shown all the comparisons containing loci with significantly higher $F_{ST}$, and one example without it). These plots were produced using the script provided by Crits-Christoph *et al*. (2020)[1].

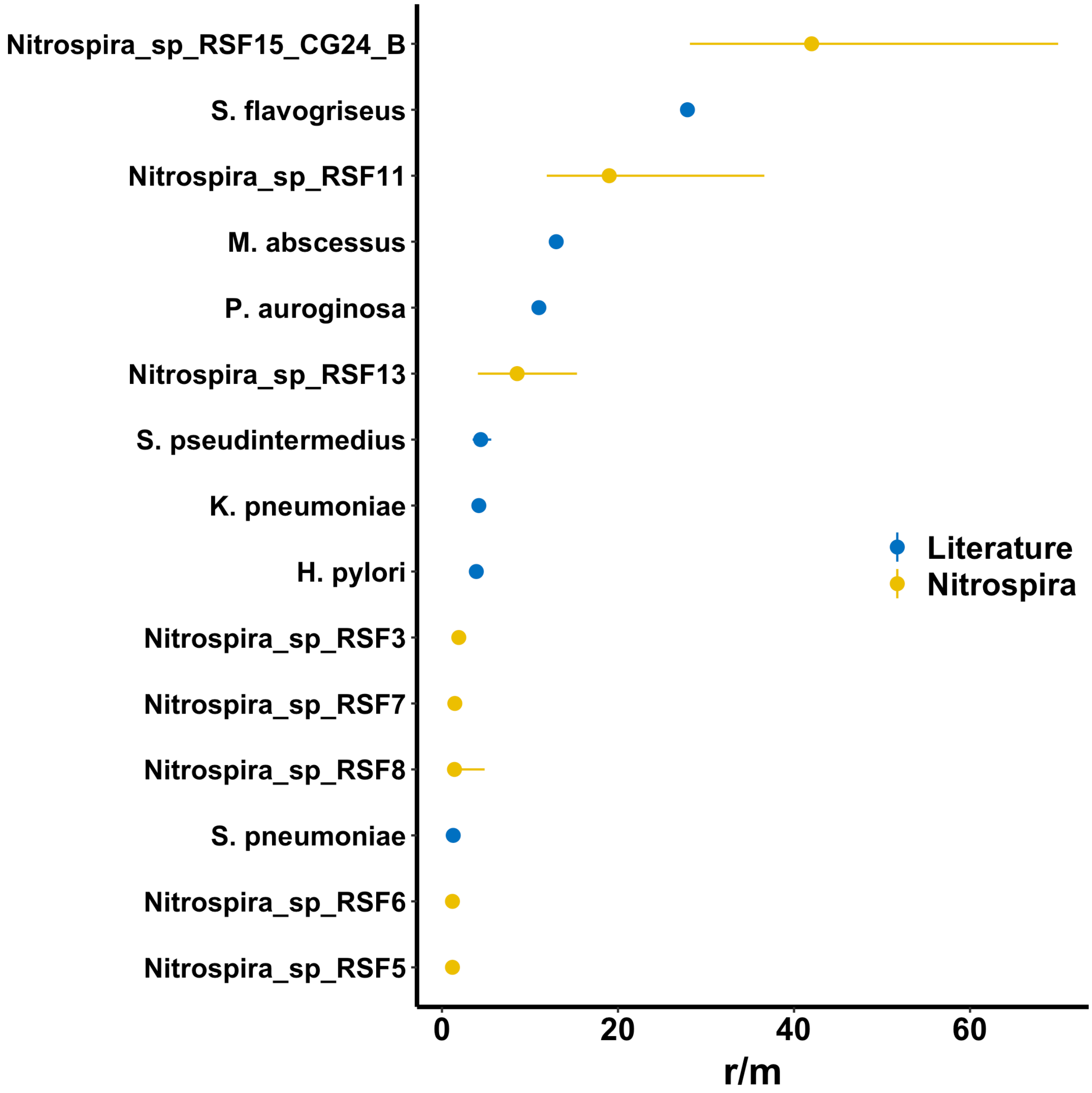


Fig. S12. Relative rates of recombination to mutation (*r/m*) calculated across the 12 waterworks for the *Nitrospira* populations (yellow) on synonymous third position codon sites, compared with previous values (blue) reported by Lin and Kussell[2], the value reported for Streptomyces flavogensis by Doroghazi and Buckley[3], and the value reported for Staphylococcus pseudintermedius by Smith et al. (2020)[4]. Error bars represent the 95% confidence interval across 1000 bootstraps.

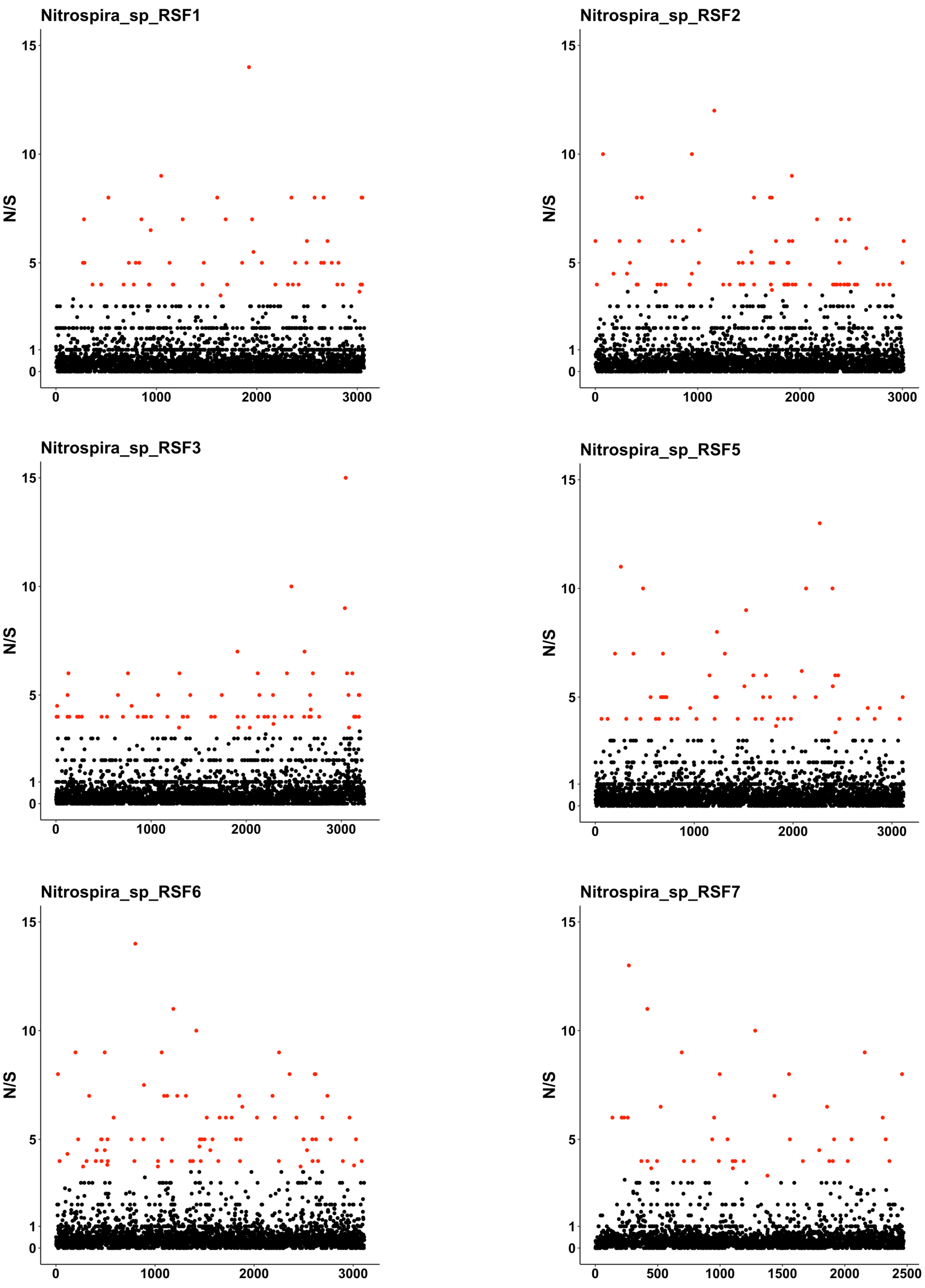

Nitrospira_sp_RSF1
Nitrospira_sp_RSF2
Nitrospira_sp_RSF3
Nitrospira_sp_RSF5
Nitrospira_sp_RSF6
Nitrospira_sp_RSF7
N/S
15
10
5
1
0
0
500
1000
1500
2000
2500
3000

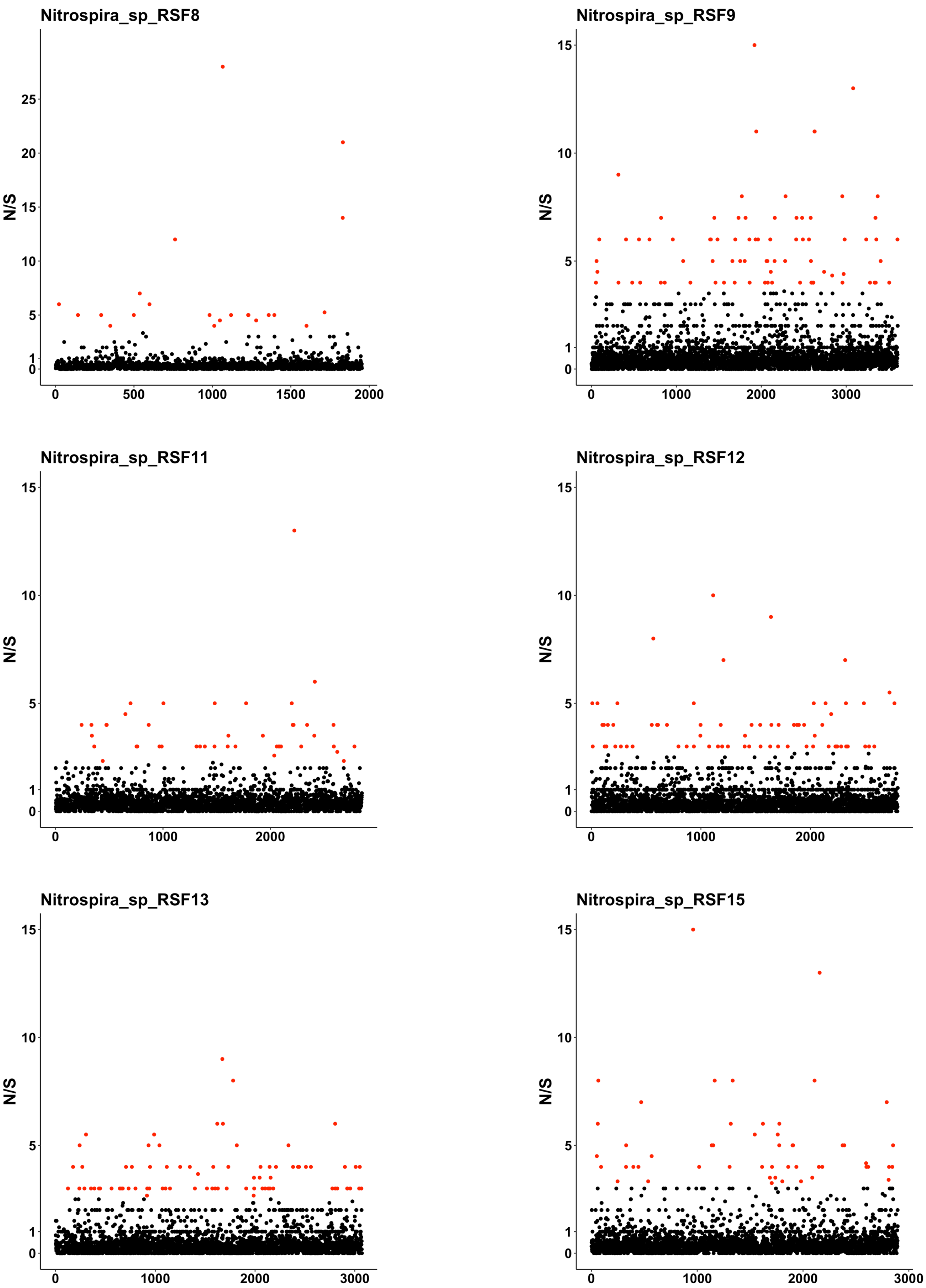
Nitrospira_sp_RSF8
N/S
Nitrospira_sp_RSF9
N/S
Nitrospira_sp_RSF11
N/S
Nitrospira_sp_RSF12
N/S
Nitrospira_sp_RSF13
N/S
Nitrospira_sp_RSF15
N/S

Fig. S13. Values of pN/pS ratios for genes across the genomes of 12 *Nitrospira* populations. Each point is a gene, and genes with significantly higher (more than three standard deviation than the mean) pN/pS value are highlighted in red.

References:


1. Crits-Christoph, A., Olm, M. R., Diamond, S., Bouma-Gregson, K. & Banfield, J. F. Soil bacterial populations are shaped by recombination and gene-specific selection across a grassland meadow. *ISME J.* 1–25 (2020). doi:10.1038/s41396-020-0655-x
2. Lin, M. & Kussell, E. Inferring bacterial recombination rates from large-scale sequencing datasets. *Nat. Methods* **16**, 199–204 (2019).
3. Doroghazi, J. R. & Buckley, D. H. Widespread homologous recombination within and between Streptomyces species. *ISME J.* **4**, 1136–1143 (2010).
4. Smith, J. T. *et al.* Population genomics of Staphylococcus pseudintermedius in companion animals in the United States. *Commun. Biol.* **3**, 282 (2020).
